# Massively parallel quantification of mutational impact on IAPP amyloid formation

**DOI:** 10.1101/2025.05.19.654797

**Authors:** Marta Badia, Cristina Batlle, Benedetta Bolognesi

## Abstract

Amyloid fibrils formed by the islet amyloid polypeptide (IAPP) cause pancreatic beta-cell damage, resulting in reduced insulin secretion and Type 2 diabetes (T2D). Variations in the primary amino acid sequence of IAPP can influence its aggregation rate and animals expressing IAPP variants that do not form amyloids, do not develop T2D. Conversely, specific single amino-acid changes in IAPP are enough to accelerate its aggregation rate. Understanding how mutations impact IAPP aggregation can help gain mechanistic understanding into the process of pathogenic amyloid formation of this peptide and preventively identify mutations that may contribute to the risk of developing T2D. Here, we employ deep mutational scanning to measure the ability to nucleate amyloids for 1663 IAPP variants, including substitutions, insertions, truncations and deletions and identify variants that increase amyloid formation in all mutation classes. Our results point at a continuous stretch of residues (15-32) which likely is structured in IAPP amyloids and that matches the core of the early aggregated species formed by IAPP *in vitro*. Inside this region, mutations have a more drastic effect in the 21-27 NNFGAIL segment, suggesting tighter structural constraints for this stretch in IAPP amyloids. Finally, by comparing this mutational atlas to that of another amyloid, Amyloid beta (Aβ42), the peptide that aggregates in Alzheimer’s Disease, we find that the effects of mutations that slow down nucleation correlate between the two amyloids, but that when it comes to mutations that accelerate nucleation one single amyloid dataset cannot be used to predict mutational effects in the other.

## Introduction

Amyloids formed by the islet amyloid polypeptide hormone (IAPP) are among the first ever amyloids observed and extracted from human tissue, more than 120 years ago^1^, decades before evidence of amyloids was observed in dementia brains.

It is now established that IAPP amyloids cause pancreatic beta-cell damage, leading to a decline in insulin secretion and Type-II diabetes (T2D)^2^, a debilitating condition which affects over 100 million patients worldwide^3^. While IAPP is highly conserved among mammals, IAPP variants that differ in less than a handful of positions from the human sequence don’t form amyloids and animals expressing these sequences, such as bear or mouse, do not develop diabetes^4,5^. It is however enough to express human IAPP in a mouse model, to cause the formation of pancreatic amyloids and diabetes to occur^6,7^. Pramlintide, a IAPP analogue that does not form amyloids, is currently approved as T2D treatment, with a need for the design of alternative IAPP analogues with improved solubility^8^.

Early aggregates formed by IAPP have been shown to be more damaging for beta-cells^2,9–11^, highlighting the need to understand and control amyloid nucleation, the initial event and rate limiting step of the amyloid reaction^12^. The challenges of quantifying amyloid nucleation rates for multiple variants in one unique set-up has so far partially limited our understanding of the sequence-nucleation relationship for IAPP^13^. It is currently unknown which mutations could increase IAPP aggregation rate and therefore it is difficult to preventively know if novel variants found in the population can contribute to increasing the risk of developing T2D.

Several studies have proposed an analogy between IAPP and another extracellular peptide, Amyloid Beta (Aβ42), the main component of the plaques that deposit in Alzheimer’s disease (AD) brains^14^. Sequence alignment of the two proteins shows 56% sequence similarity, with 9 positions (24%) having identical amino acids. Four of these identical amino acids are located in the previously described core of the fibrils formed *in vitro* by both proteins (NxGAI, positions 22, 24-26 in IAPP and 27, 29-31 in Aβ42). Different structural polymorphs have been described for fibrils formed by the two peptides *in vitro* or extracted from human tissue (8 for IAPP^14–19^ and 10 for Aβ42^20–25^). Among these, two specific Cryo-EM structures of *in vitro* fibrils display striking structural alignment^14,16^. The two peptides have also been reported to interact and co-aggregate *in vitro* and in yeast cells^26–29^.

We reasoned that a systematic approach to probe the similarities between these two sequences that form amyloids in physiological conditions is the massively parallel quantification of mutational impact to understand whether mutations speed up or slow down the amyloid reaction in a similar way. We have previously mapped the complete mutational landscape of the Aβ42 peptide^30^ and have gathered insights into pathogenic gain of function (GOF) from systematically characterizing insertions and deletions (indels)^31^. Here, we employ a similar strategy to quantify the impact of substitutions, insertions and deletions in IAPP, for a total of 1663 variants. Besides comprehensively mapping for the first time the IAPP mutational landscape and identifying the likely structured core of the nucleating IAPP fibrils, the resulting dataset reveals that GOF, i.e. the acceleration of amyloid nucleation, can result from all different classes of mutations, highlighting the need to improve our ability to measure, understand and predict the impact of indels in human sequences.

We also systematically compare the outcome of mutations in IAPP and Aβ42 and find significant similarities between mutational effects in IAPP and Aβ42 when it comes to mutations that reduce amyloid nucleation, suggesting a common mechanism of disruption of amyloid nucleation. However, the effects of GOF mutations, that instead speed up nucleation and are more frequent outside the core amyloid region of both peptides, are distinct in the two datasets. Overall, one mutational dataset cannot be used to predict the effect of mutations in the other. This highlights the challenge of predicting pathogenic GOF from primary sequence and the need to produce similar quantitative datasets for all amyloids observed in human disease.

## Results

### Massively parallel quantification of IAPP amyloid nucleation

We synthesized a library encompassing six classes of mutations: single amino acid substitutions, insertions, and deletions in IAPP, as well as truncations, larger internal deletions, and sequence alterations resulting from polymerase slippage. We then quantified the ability of each sequence to form amyloids using a cellular assay where IAPP variants are fused to the nucleation domain of the yeast prion Sup35. Nucleation of endogenous Sup35p is required for yeast survival in the lack of adenine, so that fitness-based selection becomes possible and that variants that nucleate amyloid fibrils inside yeast cells get enriched upon selection (Fig. 1a). The relative enrichment of each variant is calculated from sequencing the variant library before and after selection, providing an estimate of the ability of each variant to nucleate amyloids (see Methods).

**Figure 1.**
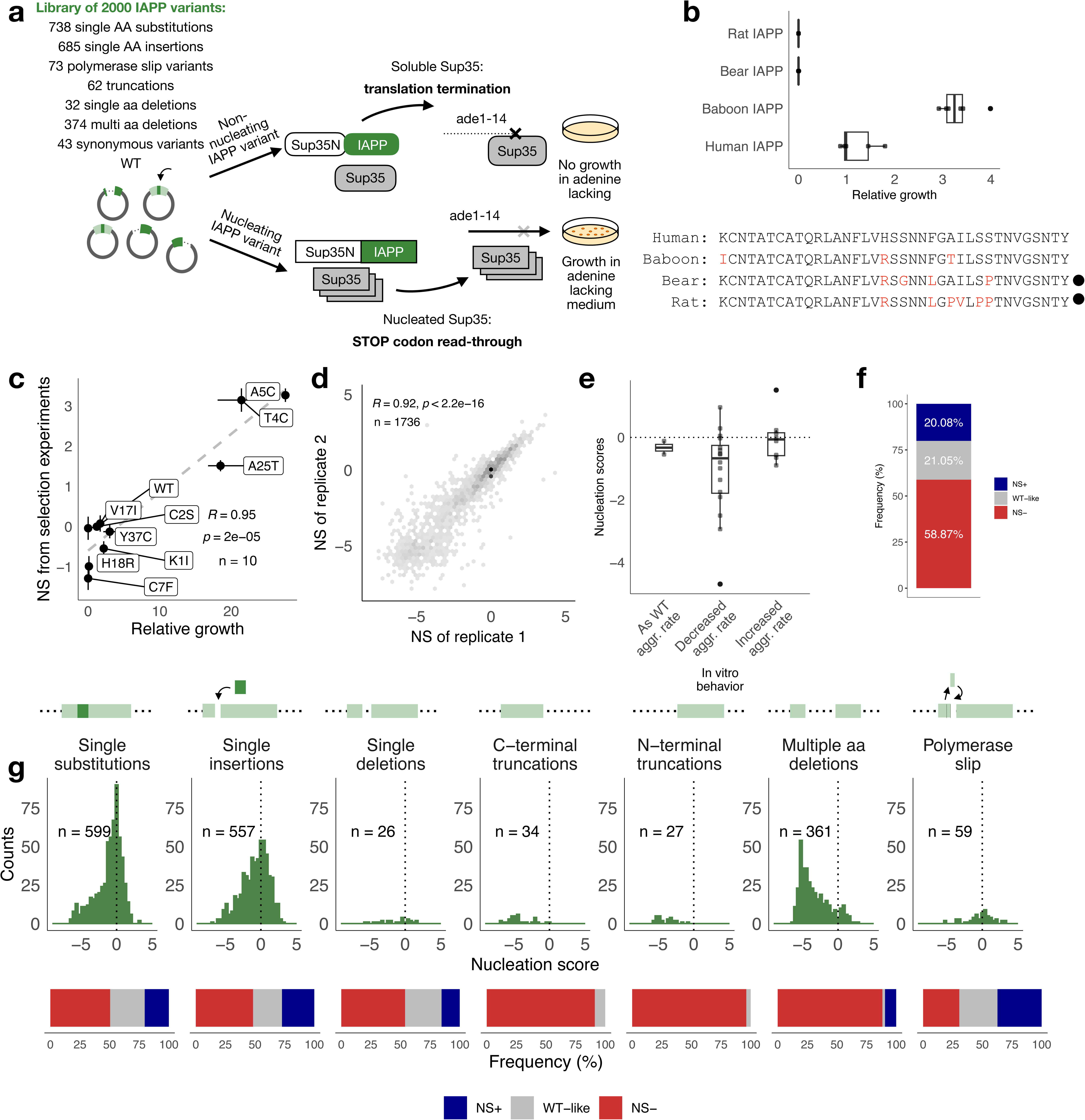
Deep mutagenesis of IAPP. **a.** Description of the IAPP library and schematics of the *in vivo* selection experiment. IAPP variants are fused to Sup35N. In a strain with a premature STOP codon in the *ade1* reporting gene, Sup35N-IAPP fusions that seed aggregation of full length Sup35p will cause STOP codon read-through and confer cells the ability to grow in adenine-lacking medium. **b.** Relative growth of cells expressing IAPP sequences from different mammals quantified as percentage of colonies grown in selective conditions (-Ade -URA) over colonies grown in non-selective conditions (-URA) (n = 4). Variants that have been reported to not nucleate in the literature are indicated with a dot. Amino acids different from human IAPP sequence are coloured in red. **c.** Correlation between nucleation scores from selection experiments and relative growth measured for individual variants in the absence of competition quantified as percentage of colonies grown in selective conditions (-Ade -URA) over colonies grown in non-selective conditions (-URA) (n = 10). Error bars indicate the standard deviation of the mean estimations. **d.** Correlation of nucleation scores between biological replicates. Pearson correlation coefficients and number of variants used in each plot are indicated in c. and d. **e.** Nucleation scores of IAPP variants grouped by their reported ability to form aggregates *in vitro*. **f.** Percentage of variants increasing (blue), decreasing (red) or having no effect (gray) on IAPP nucleation at FDR = 0.1. **g.** Distribution of nucleation scores, number of variants and fraction of variants increasing, decreasing or having no effect on IAPP nucleation at FDR = 0.1 grouped per mutation type.

As a result, we obtained a dataset reporting nucleation scores and associated error terms for 1663 IAPP variants that belong to all classes of mutations (Fig. 1g). Nucleation scores are reproducible across biological replicates (Fig. 1d and Suppl. Fig. 1) and they also correlate well with measurements of individual variants’ growth in selective conditions (Fig. 1c). By comparing nucleation scores to the results of over ten previous studies where the kinetics of amyloid formation of IAPP variants (n = 26) were followed *in vitro* by Thioflavin-T fluorescence, we show that nucleation scores accurately capture the direction of the mutational effects in speeding up or slowing down the rate of amyloid formation (Fig. 1e, Suppl. Table S1). Our assay also captures the different amyloid propensity of IAPP animal variants that have previously been characterized to form or not form aggregates *in vitro* and *in vivo* (Fig. 1b). An advantage of this deep mutational scanning approach is obtaining quantitative estimates for all IAPP variants in one unique experimental set-up, overcoming some of the challenges involved in quantifying *in vitro* rates for more than a handful of variants in identical conditions. For comparison, at most four variants were studied in parallel in previous studies, and quantitative estimates of rate constant were only obtained for WT and S20G^13^.

### Distribution of mutational effects across classes of mutations

The distribution of nucleation scores (NS) in each class of mutations reveals that mutations in IAPP are overall more likely to decrease the ability of the peptide to nucleate amyloids with 59% of mutations slowing down aggregation (NS-), while 21% of mutations behave similarly to WT IAPP and only 20% speed up its aggregation (NS+) (Fig. 1f). In those classes of mutations, where changes involve the removal of more than one amino acid such as multiple amino acid deletions and truncations from the N-terminus and the C-terminus, an even larger percentage of variants reduce nucleation (Fig. 1g, 88.3% of NS-multiple aa deletions, 96% of NS- N-terminal truncations, and 91% of NS-C-terminal truncations).

### Single amino acid substitutions highlight a structured core between residues 15 and 32

Substitutions to proline and glycine (Fig. 2b) reduce (or maintain) nucleation in a continuous stretch from residue 15 to 32, suggesting this region is the one getting structured in the amyloid fibrils nucleated by IAPP and where side-chain constraints are tighter. This region matches the structured region in all fibrils formed by IAPP *in vitro* as well as in those seeded by samples extracted from T2D patients. Within this region, we find the lowest nucleation scores in the NNFGAIL stretch (Fig. 2b), which has been shown to be essential for IAPP aggregation *in vitro* and one of the shortest peptides known to form amyloids *in vitro*^14,32,33^. Mutations of Ala 25 represent the only exception in this window, in line with *in vitro* evidence demonstrating that substitutions of Ala 25 can maintain nucleation, including the A25P variant which alone or in combination with S29P still forms amyloid fibrils^5^. The impact of mutations in the NNFGAIL stretch appears to be dominant: double mutants containing the A25T mutation (NS+) feature increased aggregation compared to the other single mutant, while the opposite happens in double mutants containing F23L (NS-) where amyloid formation is reduced relative to the other single substitutions (Fig. 2c).

**Figure 2.**
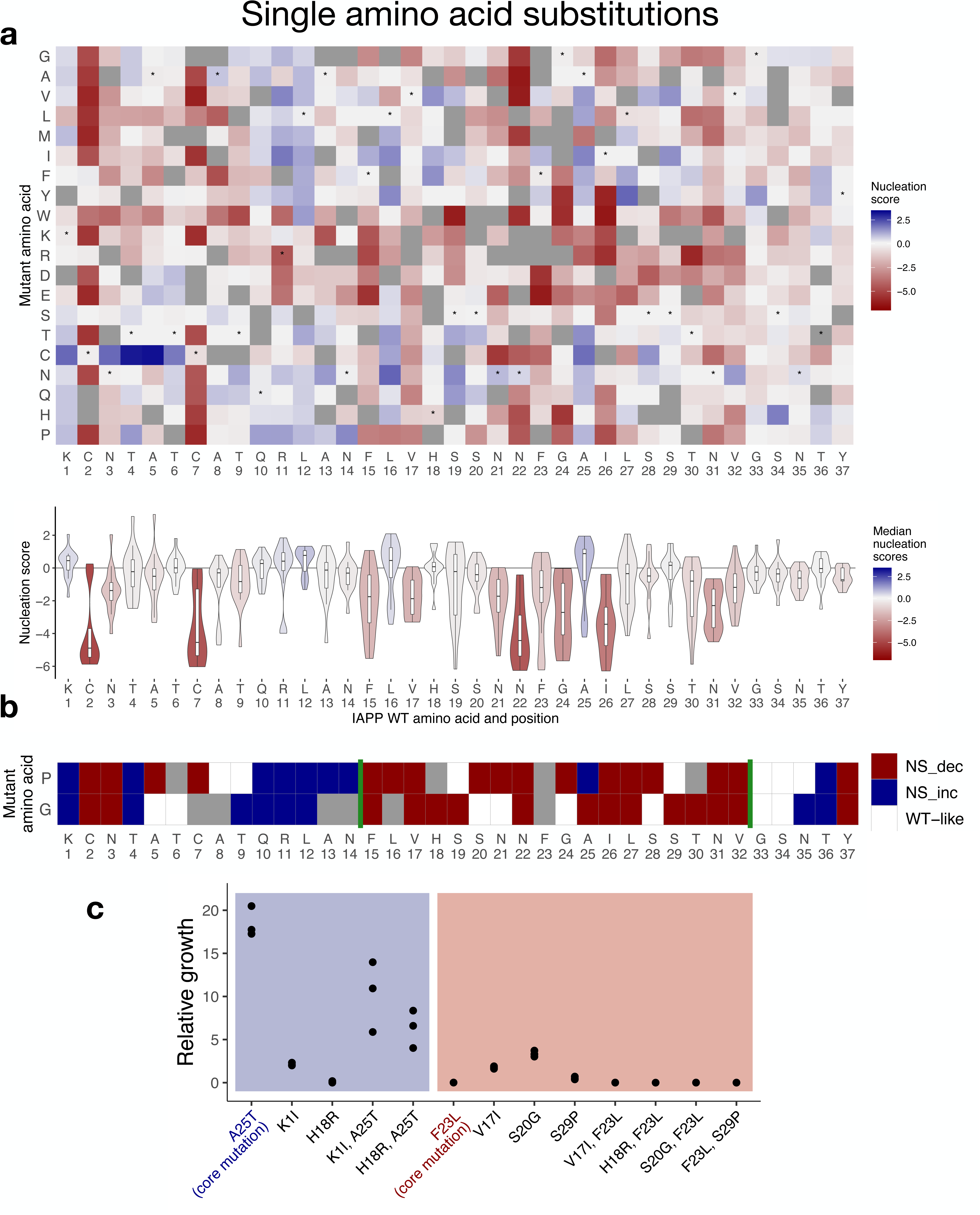
Mutational effects of IAPP amino acid substitutions. **a.** (top) Heatmap of nucleation scores of amino acid substitutions. x-axis indicates the IAPP WT sequence and the y-axis indicates the mutant amino acid. Variants not present in the library are represented in gray. Synonymous substitutions are indicated with “*”. (bottom) Distribution of the nucleation scores per position. Violin plots are colored by the mean of the nucleation score per position. **b.** FDR categories of nucleation scores for substitutions to proline and glycine along the IAPP sequence (FDR = 0.1). Missing substitutions are coloured in gray. Green lines indicate a continuous stretch of 18 aa where mutations either to proline and glycine (or both) decrease nucleation.

We also find that mutating Cys residues at positions 2 and 7 drastically reduces nucleation and that instead mutations to Cys at the first 6 positions speed up the rate of amyloid nucleation, with T4C and A5C scoring as the strongest nucleators among all IAPP substitutions (Fig. 2a).

While this fusion peptide may not recapitulate all of the modifications that *in vivo* characterize biologically active IAPP, such as C-terminal amidation and formation of the disulfide bond between Cys 2 and Cys 7, the mutational landscape of IAPP substitutions presented here identifies a likely structured region (15-32) in IAPP amyloids as well as an inner stretch of core residues (21-27) that are essential for amyloid nucleation, both of which are in line with results obtained *in vitro* with unfused oxidized peptide, overall suggesting the SupN-IAPP fusion employed in the selection assay can accurately capture the essence of IAPP nucleation.

### Mutational impact is consistent with a P-fold subunit arrangement

There is currently no high-resolution structure of IAPP fibrils from human tissue, but four structures of fibrils formed *in vitro* by recombinant or synthetic IAPP have been resolved by Cryo-EM^14–16^. In all of these, the first 11 or 12 N-terminal residues are unresolved, suggesting that, regardless of the state of the disulfide, this region is either disordered or displays structure heterogeneity in the mature fibril arrangement^34^. In fibrils seeded by patient extracts, only residues 1-5 are unresolved, but Cys 7 is not engaging in a disulphide^17^. Overall, we find 58 GOF substitutions of residues 1-12 that speed up amyloid formation of IAPP (Suppl. Fig. 2a).

We evaluated the match between mutational impact and the structural arrangement of IAPP fibrils by correlating average nucleation score per residue to the extent to which side chains are buried in each IAPP fibril structure (accessible surface area, ASA). Based on a previous mutational scan of Aβ42^31^, the expectation is that - if our assay is tracking the aggregation of IAPP into structures that are similar to those reported so far - then mutating those side chains buried in the core of amyloid fibrils should slow down nucleation the most. We find a stretch of residues (aa 21-27, with the exception of Ala 25) where more than 50% of single amino acid substitutions per position decrease the nucleation (Suppl. Fig. 3). These correspond to residues forming the minimal IAPP fragment required for amyloid formation, NNFGAIL^14,33^, and these positions also feature low relative ASA across all available IAPP fibril structures (Suppl. Fig. 3), as their side chains are buried in the fibril core. We note that mutations that affect the first of the two zippers of the P-fold structures have more drastic effects than those affecting the other, prioritizing residues 21-27 as those more likely to get structured in the initial events of IAPP aggregation (Fig. 3b).

**Figure 3.**
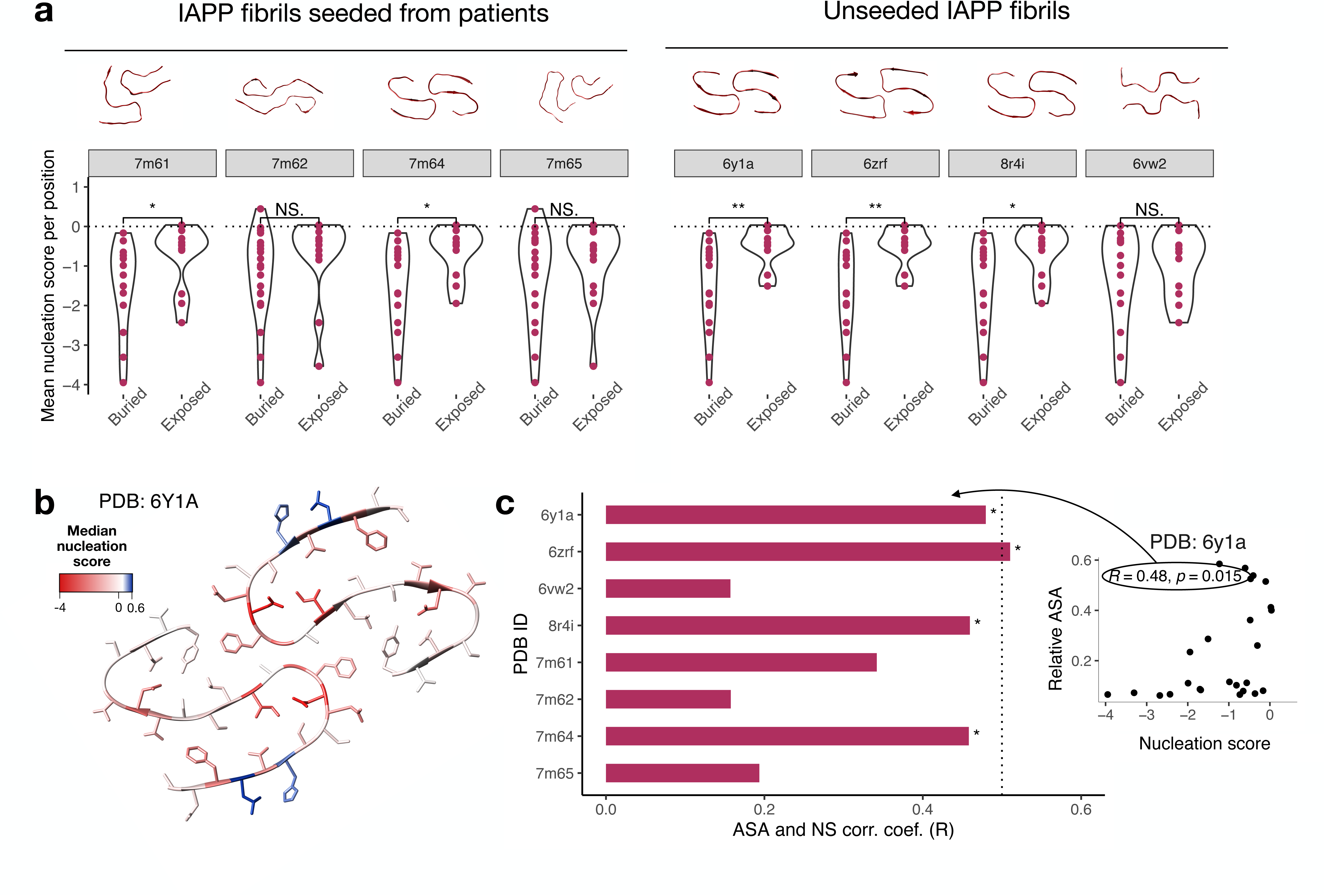
Mapping mutational impact on IAPP amyloid structures. **a.** Distribution of the averaged nucleation scores per position of single amino acid substitutions of IAPP positions classified by their exposure in all the available IAPP fibrils PDB structures. Residues with ASA > 0.25 are considered exposed. **b.** IAPP structure (PDB: 6Y1A) colored by the median mutational effect of amino acid substitutions per position. **c.** Correlation coefficients (R) of the correlations of mean nucleation scores per position and available surface area extracted from all available PDB structures. “*” indicates the correlations whose p-values < 0.05.

We also find that average nucleation scores per position are significantly different between buried and exposed residues (Fig. 3a), and that the overall correlation between nucleation score and accessible surface area (ASA) is higher for those structures where each fibril subunit adopts a P-fold (PDB: 6Y1A, 6ZRF, 8R4I, 7M64) and lower for those that consists of different folds (Fig. 3c). We note that the P-fold is also the fold of the first species identified in the early time points of the time-resolved Cryo-EM characterization of the amyloid reaction for IAPP and the IAPP S20G variant (Suppl Fig. 4, Suppl. Fig. 5).

### Half of the possible single amino acid insertions maintain or accelerate IAPP amyloid nucleation

We next evaluated the amyloid nucleation scores resulting from amino acid insertions, thus measuring the result of sequence alterations that extend the main chain. The insertions of prolines and glycines decrease nucleation in the same continuous stretch we identified measuring substitutions (15-32) (Fig. 4a, bottom panel). Within this window, all insertions after position 22, 23, 24 and 25 virtually reduce nucleation, confirming the coordinates of the essential inner core of IAPP amyloids. The N-terminus (aa 1-9) of the peptide is also sensitive to insertions (Fig. 4d). Here, the vast majority of these mutations reduce nucleation - with the exception of insertions of cysteines which, similar to substitutions to cysteines in this region, increase the rate of nucleation. (Fig. 4e)

**Figure 4.**
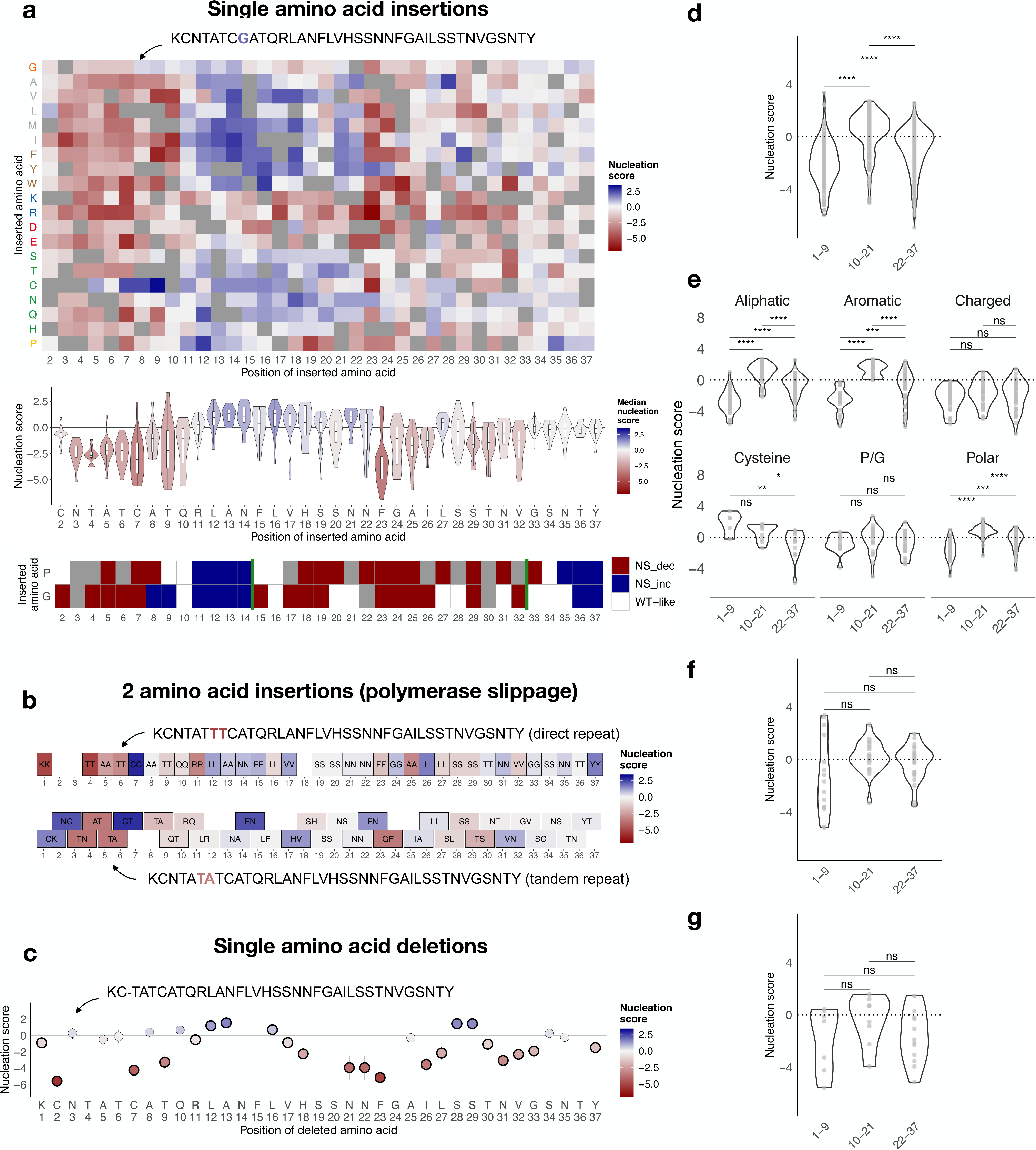
Mutational effects of IAPP amino acid insertions and deletions. **a.** (top) Heatmap of nucleation scores of 1 amino acid insertions. x-axis indicates the position of the inserted amino acid and the y-axis indicates the amino acid inserted. Variants not present are represented in gray. (center) Distribution of the nucleation scores per position. Violin plots are coloured by the mean of the nucleation score per position. (bottom) FDR categories of nucleation scores for insertions of proline and glycine across IAPP sequence (FDR = 0.1). **b.** Nucleation scores of 2-amino acid insertions resulting from polymerase slippage. x-axis indicates the position after each pair of amino acids is inserted. The inserted amino acid pair is written inside each tile. Colours represent the nucleation scores of each variant. Variants with nucleation scores significantly different from WT (FDR = 0.1) are indicated with a black square. **c.** Effect of single amino acid deletions on IAPP nucleation. x-axis indicates the position of the amino acid deletion. Nucleation score of each deletion is indicated in the y-axis and with the colour of each point. Variants with nucleation scores significantly different from WT (FDR = 0.1) are indicated with a black circle. Vertical bars represent the error. Horizontal line indicates the nucleation score of WT IAPP. **d.** Distribution of nucleation scores of all 1 amino acid insertions in the clusters obtained by hierarchical clustering of their nucleation scores (Suppl. Fig. 5). **e.** Distribution of nucleation scores of all 1 amino acid insertions separated by type of amino acid inserted. **f.** Distribution of nucleation scores of 2 amino acid insertions and **g.** single amino acid deletions. Pairwise t-test significance is indicated in **d.**, **e**., **f.** and **g**.

Systematic mapping of single amino acid insertions and hierarchical clustering of nucleation scores per position (Suppl. Fig. 6a, Suppl. Fig. 6b) also reveals a central region of the peptide, residues 10-21, where all insertions of aliphatic, aromatic and polar residues increase nucleation (Fig. 4d, Fig. 4e). What is more, between residues 10 and 14 even the insertion of prolines and glycines speeds up nucleation, suggesting the presence of an element of secondary structure able to stabilize the IAPP monomeric ensemble against amyloid nucleation. When insertions disrupt it, nucleation becomes more likely, suggesting this region could represent a potential target for small molecule binders able to modulate IAPP nucleation. Binding of chaperones stabilizing this region has been shown to inhibit fibril formation^35^.

We have also evaluated the effect of double amino acid insertions that can result from polymerase slippage, i.e. the duplication of two amino acids of the WT sequence, or of one same amino acid, repeated twice. Overall, these insertions of two amino acids have mixed effects, with variants that increase or decrease nucleation across the whole sequence (Fig. 4b, Suppl. Fig. 7). The effect of single amino acid insertions can be used to predict the impact of double amino acid insertions at the same position, especially for those where the same amino acid is repeated (R = 0.73, p-value = 4.2e-06) (Suppl. Fig. 8a). However, effects of single insertions would be difficult to predict on the basis of the single amino acid substitutions alone (R = 0.32, p-value = <2.2e-16) (Suppl. Fig. 9a), similar to what was found for the effect of insertions on protein folding^36^.

Finally, the insertion of just one single cysteine (due to single or double amino acid insertions) is enough to increase nucleation (Suppl. Fig. 10a). Together with the observation that nucleation is also increased for sequences where mutations result in two adjacent cysteines (Suppl. Fig. 10c), which could by no means form a disulfide bond, this also points at a distinct mechanism from cysteine oxidation by which increasing the number of cysteines just in the first 10 residues of the peptide can favor IAPP aggregation.

### 14 single amino acid deletions maintain or increase nucleation rates

In parallel, we measured the impact of single and multi-AA deletions of the IAPP sequence (Fig. 4c). Their effects are more drastic when deletions cause loss of Cys 2, Cys 7, Thr 9, and of residues in the NNFGAIL stretch, with the exception of Ala 25 which, in line with the results of substitutions, seems dispensable for the formation of the inner core of IAPP amyloids. Deletions of single residues from position 11 to 16 increase or maintain nucleation, suggesting that alterations of the main chain in this region, other than insertions, can unlock faster nucleation. Finally, also the individual loss of Ser28 or Ser29, which results in two identical sequences, increases nucleation.

Most of the large multi-AA disrupt nucleation, but 22 deletions of just 2-4 amino acids, at different positions along the sequence, can increase it (Suppl. Fig. 11). Among these, we highlight those deletions that don’t reduce the number of cysteines, but rather bring the WT cysteines closer to each other, significantly speeding up nucleation (Suppl. Fig. 10b). We also observe a contrast between the effect of the loss of two serines, just before or just after the inner core NNFGAIL (S19, S20). The first double deletion (Δ19-20) speeds up the aggregation process, while the latter (Δ28-29) slows it down, in contrast with the individual loss of Ser28 and Ser29. Finally, deletion of the entire 10-14 stretch (QRLAN) and other shorter deletions in this region result in sequences that nucleate faster than WT, further suggesting the presence of a gate-keeping element of secondary structure in this region of IAPP, which in the WT sequence limits aggregation rates.

### The same types of mutations decrease nucleation in IAPP and Amyloid Beta

To assess the similarity of mutational impact in different amyloidogenic peptides we compared the IAPP nucleation landscape with that of Aβ42^31^. We find a positive correlation between the average effect of substitutions to (R = 0.81, p-value = 1.4e-5) or insertions of (R = 0.5, p-value = 0.026) the same amino acids in the two proteins (Fig. 5f and 5g) as well as when comparing the average effect of replacing the same amino acids (R = 0.69, p-value = 0.0091) (Fig. 5e). We note that these correlations are driven by values in the negative range for both peptides, while they are not significant for those mutations increasing nucleation (Suppl. Fig. 12, Suppl. Fig. 13).

**Figure 5.**
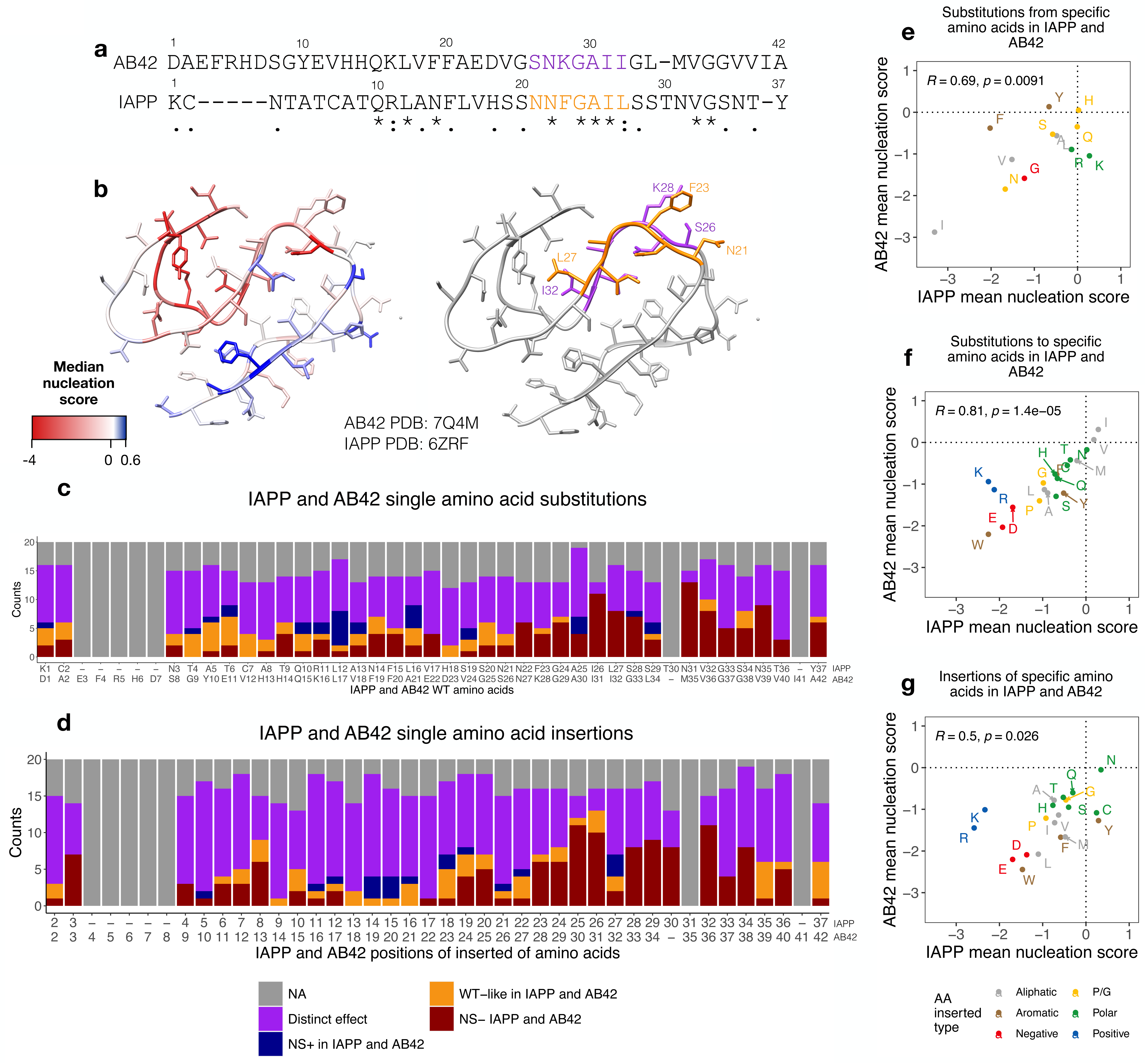
Comparison of mutational effects of IAPP and Aβ42. **a.** Sequence alignment of IAPP and Aβ42 sequence by T-COFFEE. WT positions of each protein are indicated above its sequence and gaps are indicated by “-”. Conservation scores are indicated below the alignment: “*”, “:”, “.” indicate identical amino acids, conservative changes, and semi-conservative changes, respectively. **b.** Structural superposition of IAPP (PDB: 6RZF) and Aβ42 (PDB: 7Q4M). Structures are colored by the median mutational effect of amino acid substitutions per position (left). The proposed inner core of the fibrils is highlighted for each protein (right). **c.** FDR categories (FDR = 0.1) of nucleation scores of single amino acid substitutions and **d.** insertions in IAPP and Aβ42 merged according to T-COFFEE sequence alignment. Missing variants in one or both datasets are colored in gray. x-axis indicates the WT position and sequence of IAPP and Aβ42. **e.** Correlation of the mean nucleation scores of substitutions of specific amino acids, **f.** substitutions to specific amino acids and **g**. insertions of specific amino acids in IAPP and Aβ42.

We also evaluated whether these similarities were conserved at a positional level. After aligning IAPP and Aβ42 sequences using T-COFFEE^37^, we compared the effects of 526 mutations which correspond to substitutions to the same amino acid occurring at aligned positions, with the goal of classifying mutations according to their effects on nucleation in both proteins. Out of 526 mutations, 231 (∼44%) impact the nucleation of both proteins in the same direction. Most of these mutations decrease nucleation (n = 139, NS- in IAPP and NS- in Aβ42, FDR = 0.1), and are found more frequently towards the C-terminus, where the core of the mature fibrils of both peptides is (Fig. 5c and d). In contrast, only a small number of these mutations can increase nucleation in both proteins (n = 29, NS+ in IAPP and NS+ in Aβ42, FDR = 0.1) (Fig. 5c and d), implying that, while disruption of amyloid formation proceeds in a similar way, the way the nucleation reaction can be accelerated for the two peptides is distinct.

In addition, we find that aggregation and secondary structure predictors (Zyggregator, TANGO, Camsol, s4pred) and variant effect predictors (AlphaMissense and PopEVE) perform poorly when predicting the effects of mutations on nucleation scores in both IAPP and Aβ42^31^ (Suppl. Fig. 14).

### Comparing nucleation scores to allele frequency in the population and in diabetes patients

50 IAPP substitutions have been identified in the human population, but their clinical significance remains unknown^38^. In this work, we quantified the nucleation of 48 of them: 16 NS-, 20 NS+ and 12 WT-like. (Suppl. Fig. 15, FDR = 0.1). The variant with highest allele frequency, S20G, has been extensively characterized in terms of their association to T2D and *in vitro* aggregation with overall inconclusive evidence^13,39–42^ (Suppl. Table S1). We find that, when it comes to amyloid nucleation, this sequence does not differ significantly from WT (Suppl. Fig. 2a).

We also probed whether IAPP variants for which we measure increased amyloid formation could contribute to T2D diabetes by employing the UK Biobank database which collects whole-genome sequencing and health parameters for 500,000 individuals. We retrieved 27 IAPP variants from the database^43^: 6 NS+, 8 NS- and 13 WT-like. We find none of the NS-variants in individuals with a diabetes diagnosis or with glycated hemoglobin (HbA1c) levels above 48 mmol/mol, a standard metric to diagnose and monitor diabetes^44^ (Fig. 6a and 6b). We employed burden testing to perform rare variant analysis (SKAT analysis and odds ratio) and find that individuals with NS+ mutations have higher odds ratio of being diagnosed with diabetes compared to those with other types of mutations, albeit with large confidence intervals (OR = 4.25, 95% confidence interval 1.1-12.28, p = 0.037) (Fig. 6c, Suppl. Table S4). The same tendency, although the association is not significant (OR = 4.97, 95% confidence interval 0.98-16.07, p = 0.052), is observed in individuals with high HbA1c levels (Fig. 6d, Suppl. Table S4).

**Figure 6.**
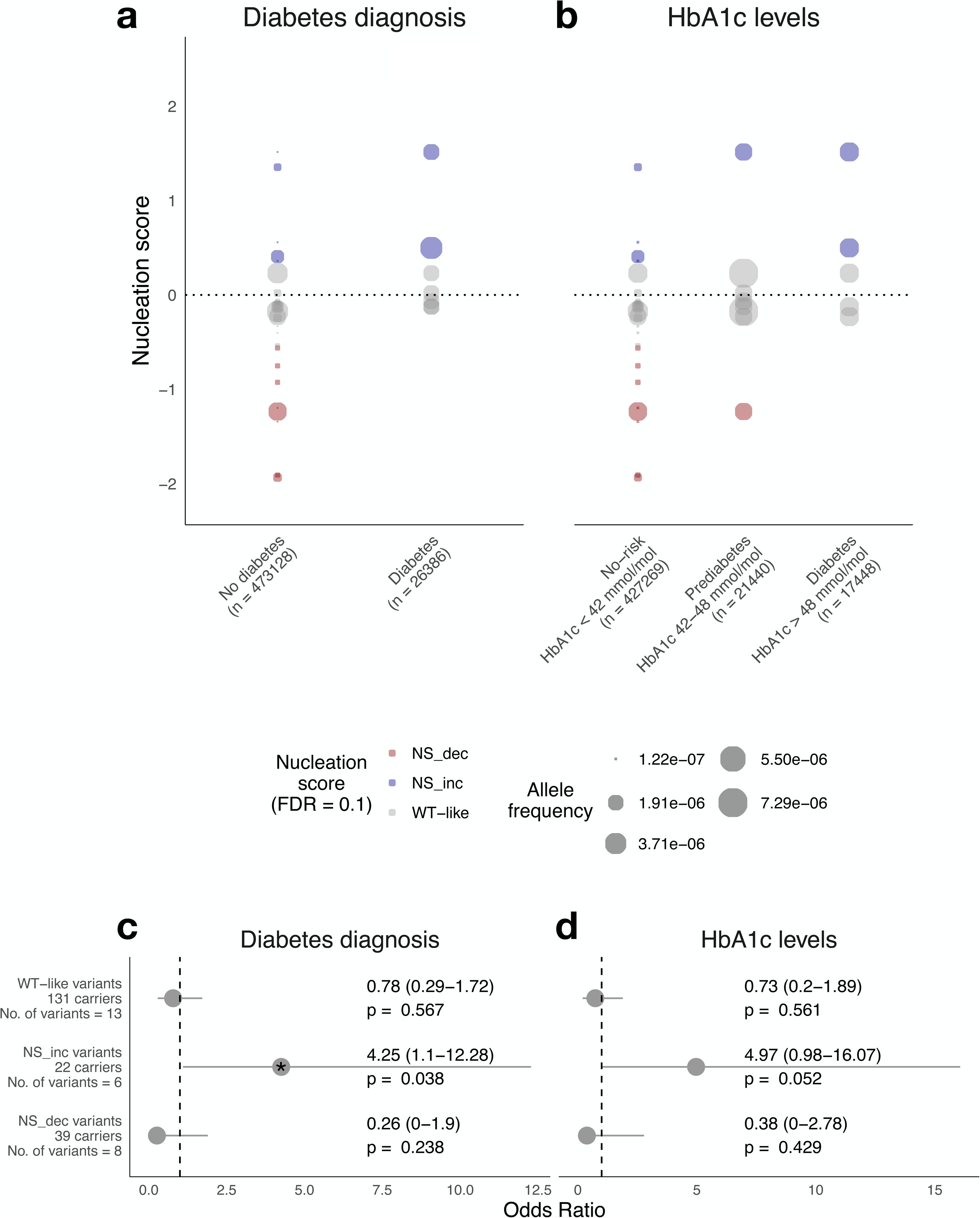
Effect of IAPP variants on T2D diabetes risk. **a.** Comparison of nucleation scores (y-axis) and allele frequencies of IAPP variants in participants grouped by diabetes diagnosis and **b.** by HbA1c levels. Variants are coloured according to their effect on IAPP nucleation (NS+, WT-like, NS-, FDR = 0.1). Point size represents allele frequency of each IAPP variant in each group. **c.** Forest plots showing odds ratio, 95% confidence interval and p-values for each IAPP variant group for diabetes diagnosis and **d.** for HbA1c levels. Number of variants and carriers in each group are indicated. “*” denotes statistically significant odds ratio.

## Discussion

Here, we present the first comprehensive mutational landscape of IAPP amyloid nucleation, covering 599 single amino acid substitutions, 557 single amino acid insertions, 59 2-amino acid insertions resulting from polymerase slippage and 448 single and multi amino acid deletions. This atlas reveals 334 GOF mutations that are able to speed up the aggregation of IAPP. These fast nucleators result from mutations of all types, including insertions, deletions and substitutions out of the structured core of IAPP amyloids which would have been hard to predict using currently available aggregation predictors or variant effect predictors.

It would have been difficult to predict these GOF variants even on the basis of the complete mutational landscape of Aβ42, another peptide that aggregates extracellularly and that can, under specific circumstances, form fibrils that structurally align to those formed by IAPP. This suggests that simply one dataset is not enough to recapitulate GOF mutational effects in another amyloid dataset and highlights the need to generate variant effect maps for all human amyloids. Similarities between the two datasets are limited to those mutations that disrupt amyloid nucleation, where changes in nucleation caused by the same types of mutations correlate strongly between datasets. Second, in both Aβ42 and IAPP, two interfaces build the structured core of mature fibrils, where strands face each other in a steric zipper. Our results reveal that, for both peptides, mutations are more disruptive in one of the two interfaces.

GOF mutations are found more frequently outside of these core regions for both peptides, including those parts of the sequence which are not even resolved in the final structures of mature amyloid fibrils (1-12 for Aβ42 and 1-11 for IAPP). More specifically, in IAPP, we find that 35,6% of single amino acid substitutions and insertions up to residue 15 are GOF, and identify a region (aa 10-21) where 110 out of 214 (51,4%) insertions speed up nucleation, hinting at the presence of a secondary structure element that in the WT could protect against aggregation by stabilizing the IAPP monomeric ensemble.

While preventively quantifying amyloid nucleation for 1663 IAPP variants in one unique set-up, our study comes with a series of limitations. Given that nucleation takes place in the yeast cytoplasm, it is likely we are only tracking mutational impact on the reduced version of the peptide. The N-terminal cysteine residues are not part of the published structure of IAPP fibrils and several *in vitro* studies have shown that IAPP forms amyloids *in vitro* both in its reduced and oxidized state^45^. However, our results revealed that mutating cysteines and mutation to cysteines have drastic effects, decreasing and increasing nucleation respectively; we cannot exclude that in this specific experimental set-up cysteines favor nucleation by other mechanisms^46^ which include multimerization via metal chelation. However, there are different lines of evidence suggesting our method accurately captures IAPP amyloid nucleation: 1) nucleation scores discriminate significant differences amongst IAPP sequences from different animals, 2) nucleation scores match with most previous *in vitro* experiments performed in different conditions on unfused peptides and 3) the pattern of mutational impact identifies a core region which corresponds to the minimal IAPP fragment essential for amyloid formation *in vitro*.

The lack of IAPP variants clearly classified and reported on Clinvar and the limited number of IAPP variants currently available in the UK Biobank make it challenging to validate this dataset with human genetics in a conclusive manner. While we find that individuals carrying variants that increase nucleation have slightly higher odds of being diagnosed with T2D, the genetic validation of the dataset, unlocking its use to contribute to clinical variant classification, will have to wait for additional data from sequencing of new T2D patients. However, the preventive quantification of amyloid nucleation presented here can contribute to the classification of these variants whenever they are encountered in the clinic or in healthy individuals. There is also a critical need for the design of soluble IAPP analogs. The dataset presented here can help guide the design of IAPP variants that do not aggregate into amyloids but that are still active and more bioavailable than the currently employed pramlintide.

Finally, this work highlights the importance of developing selection assays that, combined to deep mutagenesis, allow the characterization of novel pathogenic gain of function variants, which are particularly hard to predict and to characterize, yet are important drivers of human disease.

**Supplementary Figure 1.**
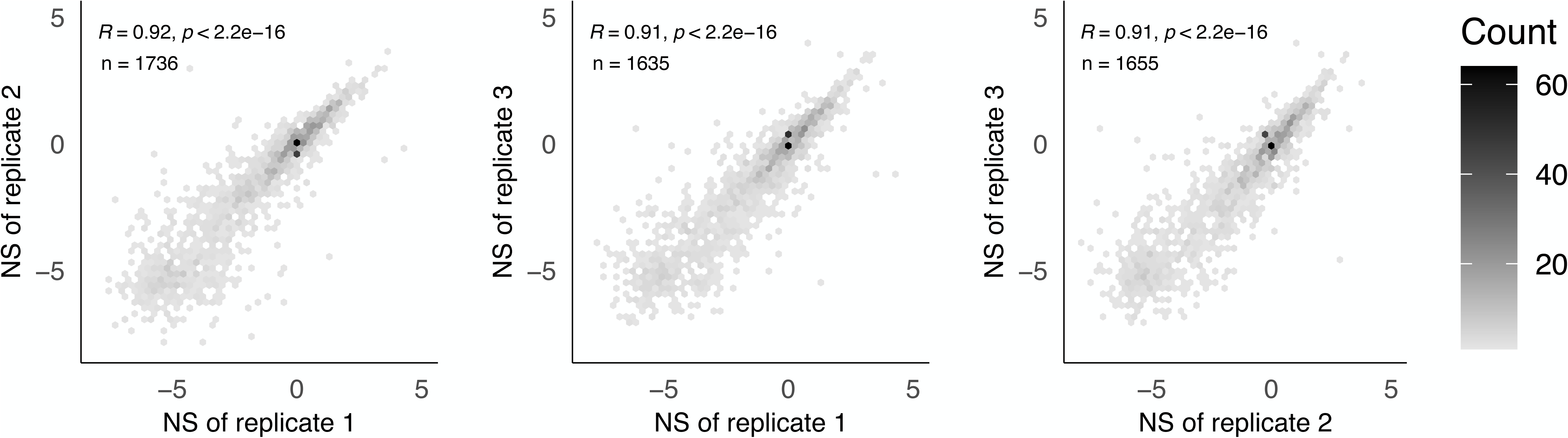
Reproducibility of the nucleation assay. Correlation of nucleation scores for 3 biological replicates of the selection assay.

**Supplementary Figure 2.**
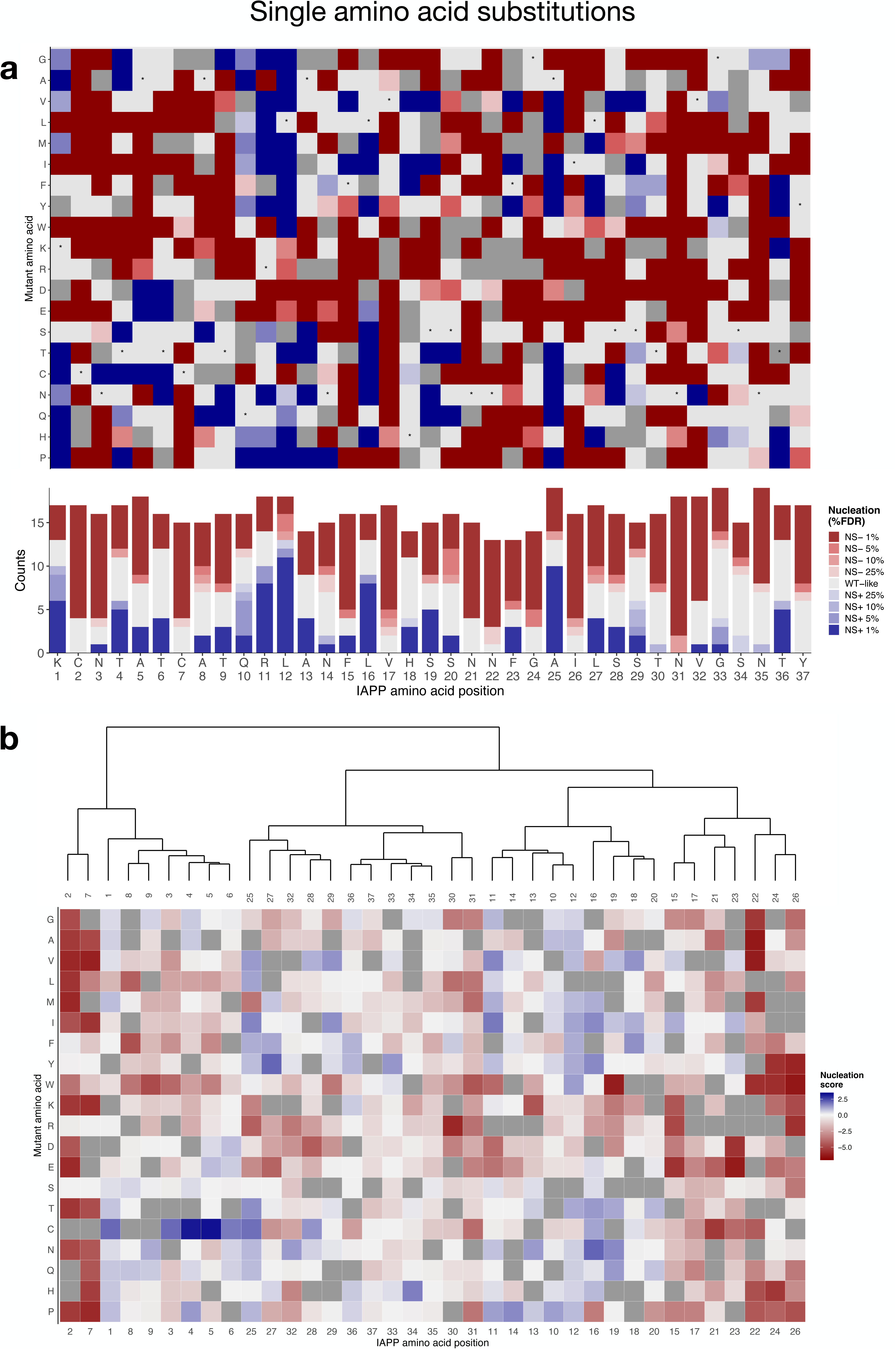
Mutational effects of IAPP single amino acid substitutions. **a.** (top) Heatmap of nucleation scores FDR = 0.1 categories for single amino acid substitutions. x-axis indicated IAPP WT position and the y-axis indicated the amino acid mutated. Variants not present in the library are represented in gray. Synonymous mutants are indicated with ‘*’. (bottom) Frequency of single amino acid substitutions increasing or decreasing nucleation for each IAPP position. **b.** Hierarchical clustering IAPP positions based on nucleation scores of single amino acid substitutions.

**Supplementary Figure 3.**
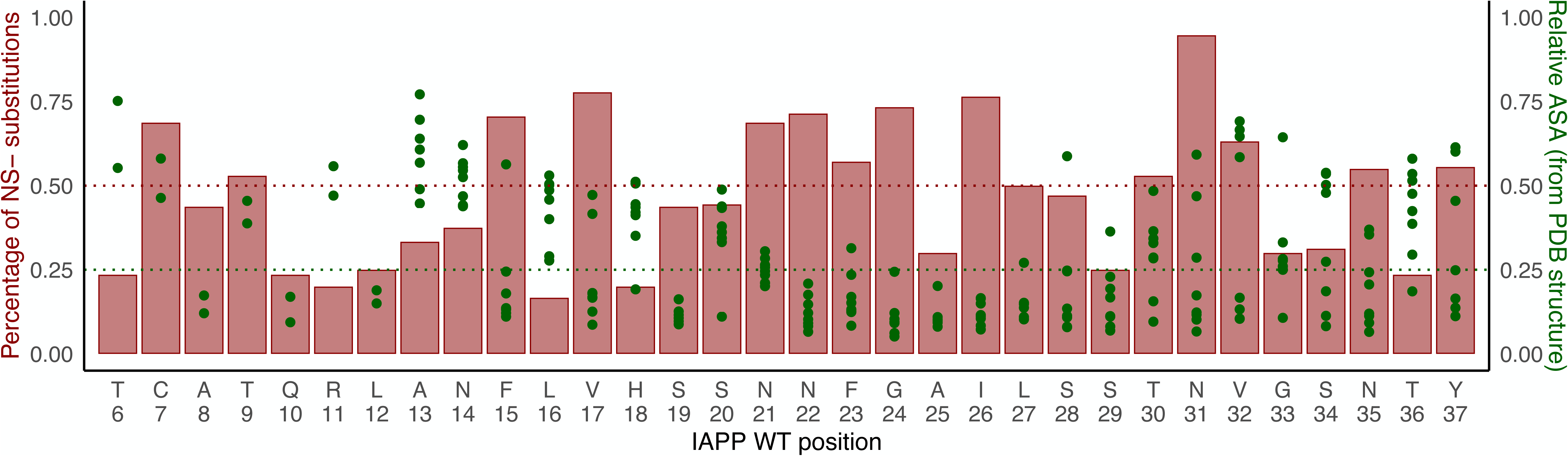
Comparison of ASA and mutations decreasing nucleation per position. The bar plot and the left y-axis indicate the percentage of single amino acid substitutions that decrease nucleation (NS-, FDR = 0.1) per each IAPP residue. The red dotted line indicates a threshold of 50%. Green dots and the right y-axis indicate the relative ASA per position of each of the 8 IAPP fibril structures available from PDB. Points below the green dotted line (relative ASA = 0.25) correspond to residues considered buried in a specific PDB structure.

**Supplementary Figure 4.**
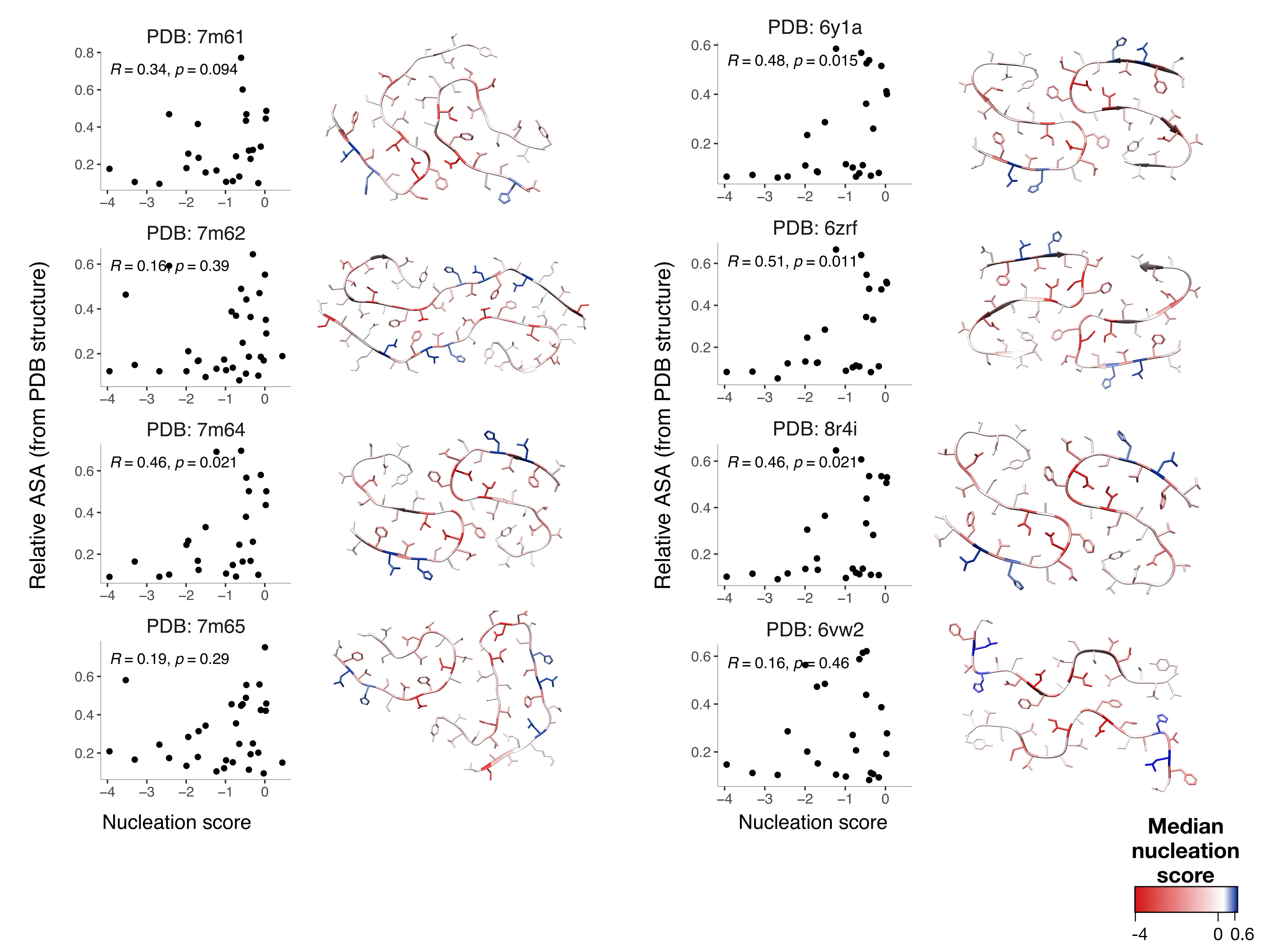
Mutational effect of IAPP substitutions on all reported IAPP WT PDB fibril structures. Correlation of mean nucleation scores of substitutions per position and available surface area extracted from all PDB reported IAPP WT structures. Pearson correlation coefficients and p-values are indicated. IAPP WT fibril structures are colored by the median mutational effect of amino acid substitutions per position.

**Supplementary Figure 5.**
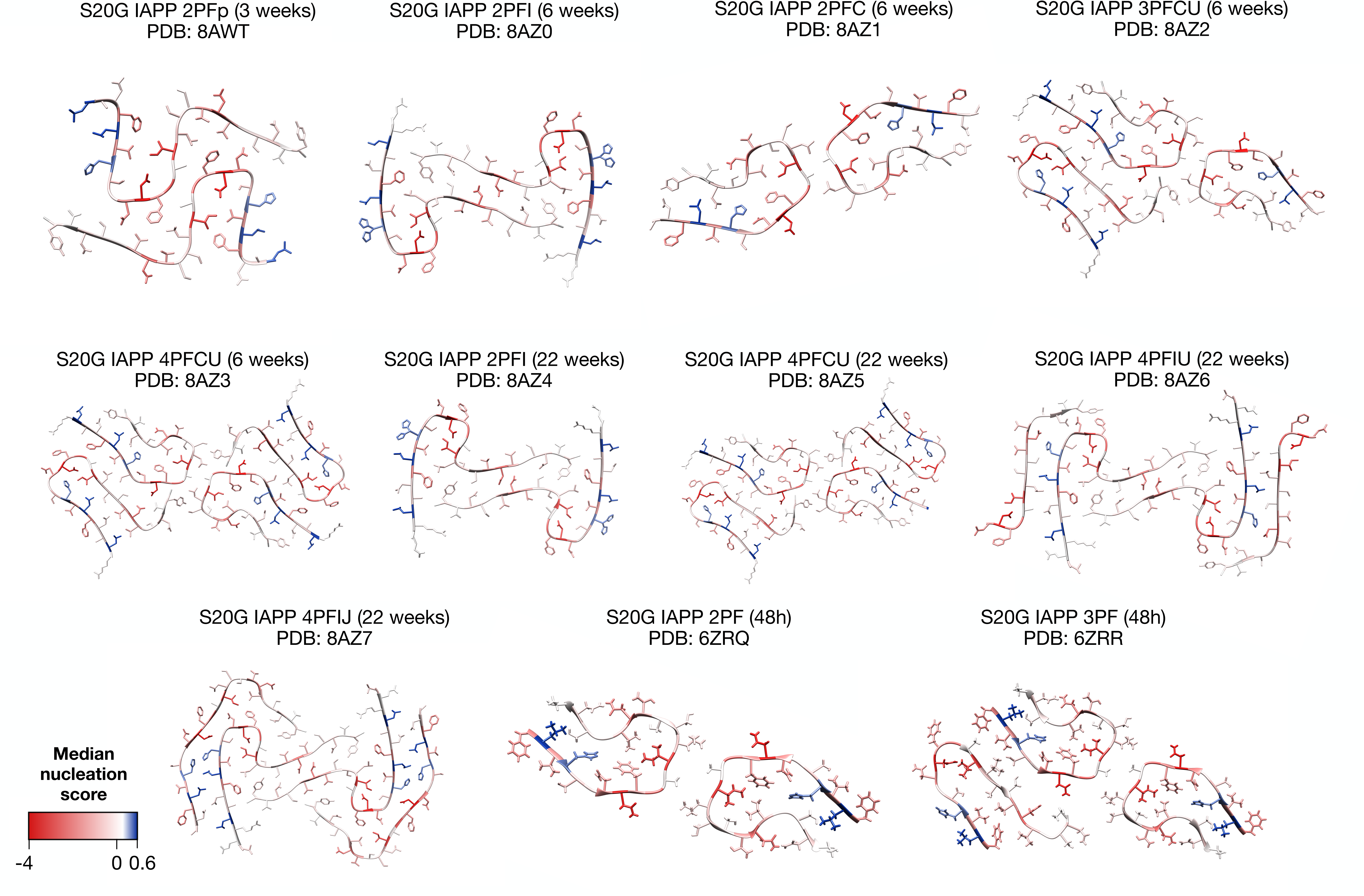
Mutational effect of IAPP substitutions on all reported IAPP S20G PDB fibril structures. IAPP S20G fibril structures from the PDB colored by the mean mutational effect of amino acid substitutions per position.

**Supplementary Figure 6.**
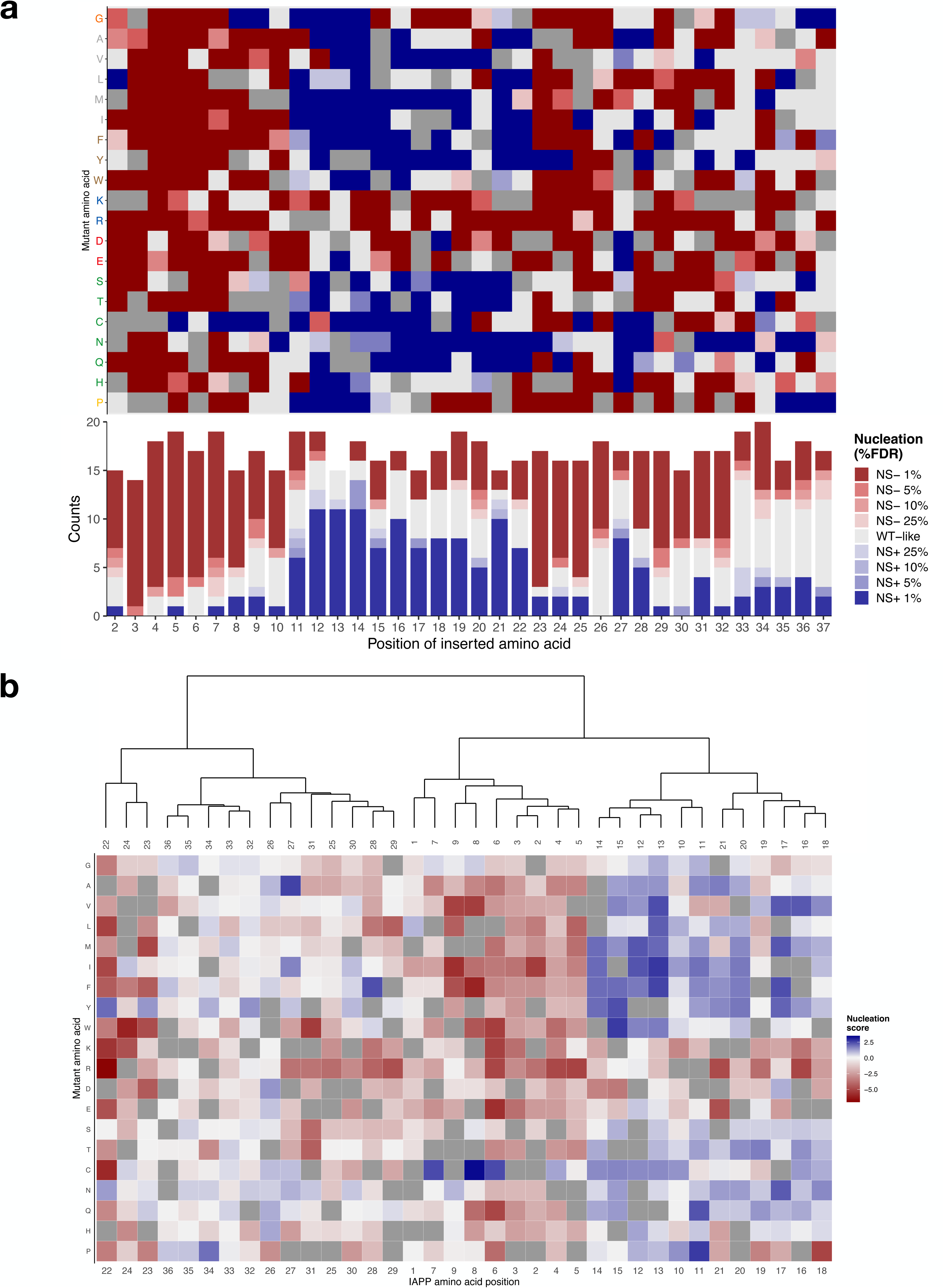
Mutational effects of IAPP single amino acid insertions. **a.** (top) Heatmap of nucleation scores FDR = 0.1 categories for single amino acid insertions. x-axis indicates the position of the inserted amino acid and the y-axis indicates the amino acid inserted. Variants not present are represented in gray. (bottom) Frequency of single amino acid insertions increasing or decreasing nucleation for each position. **b.** Hierarchical clustering IAPP positions based on nucleation scores of single amino acid insertions.

**Supplementary Figure 7.**
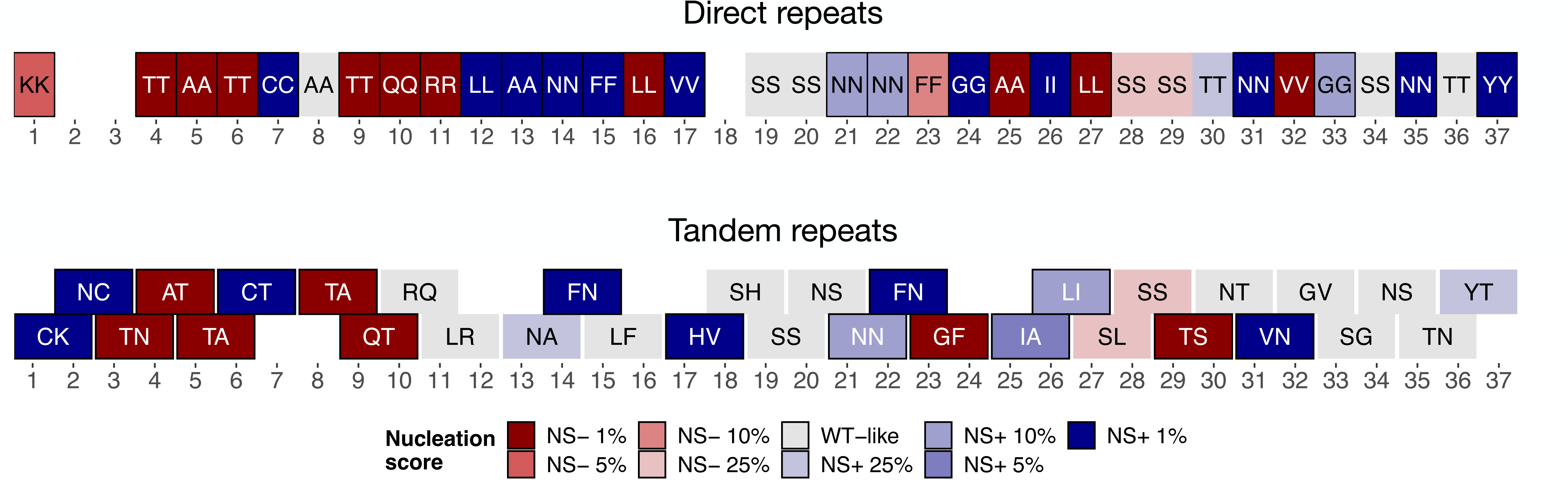
Mutational effect of IAPP variants resulting from polymerase slippage. Polymerase slippage generated variants are coloured by their impact on increasing and decreasing nucleation at different FDRs. x-axis indicates the position after each pair of amino acids is inserted. The inserted amino acid pair is specified inside the squares.

**Supplementary Figure 8.**
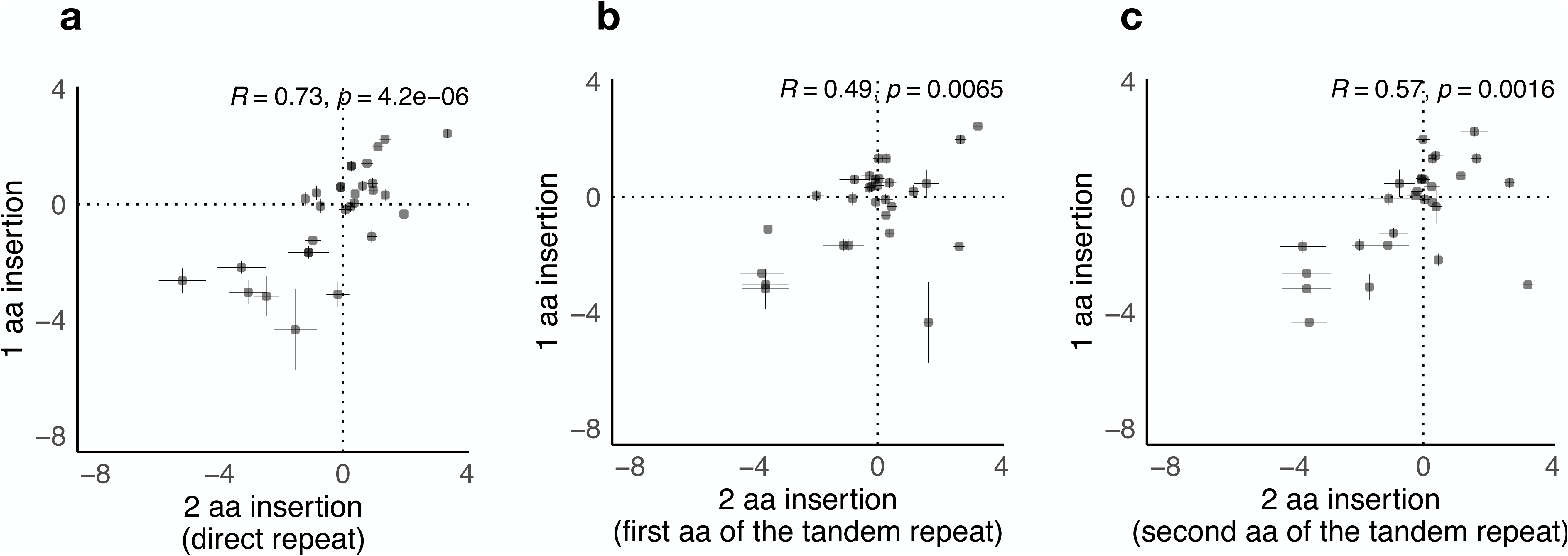
Comparison between the mutational effects of single and double amino acid insertions in IAPP. **a.** Correlation of nucleation scores resulting from single amino acid insertions and double amino acid coming from direct repeats (same amino acid repeated twice) in the same position. **b.** Correlation of nucleation scores of single amino acid insertions with the first or the **c.** second amino acid of double amino acid insertions of tandem repeats (duplication of a pair of amino acids of the WT sequence). Pearson correlations are indicated inside each correlation plot.

**Supplementary Figure 9.**
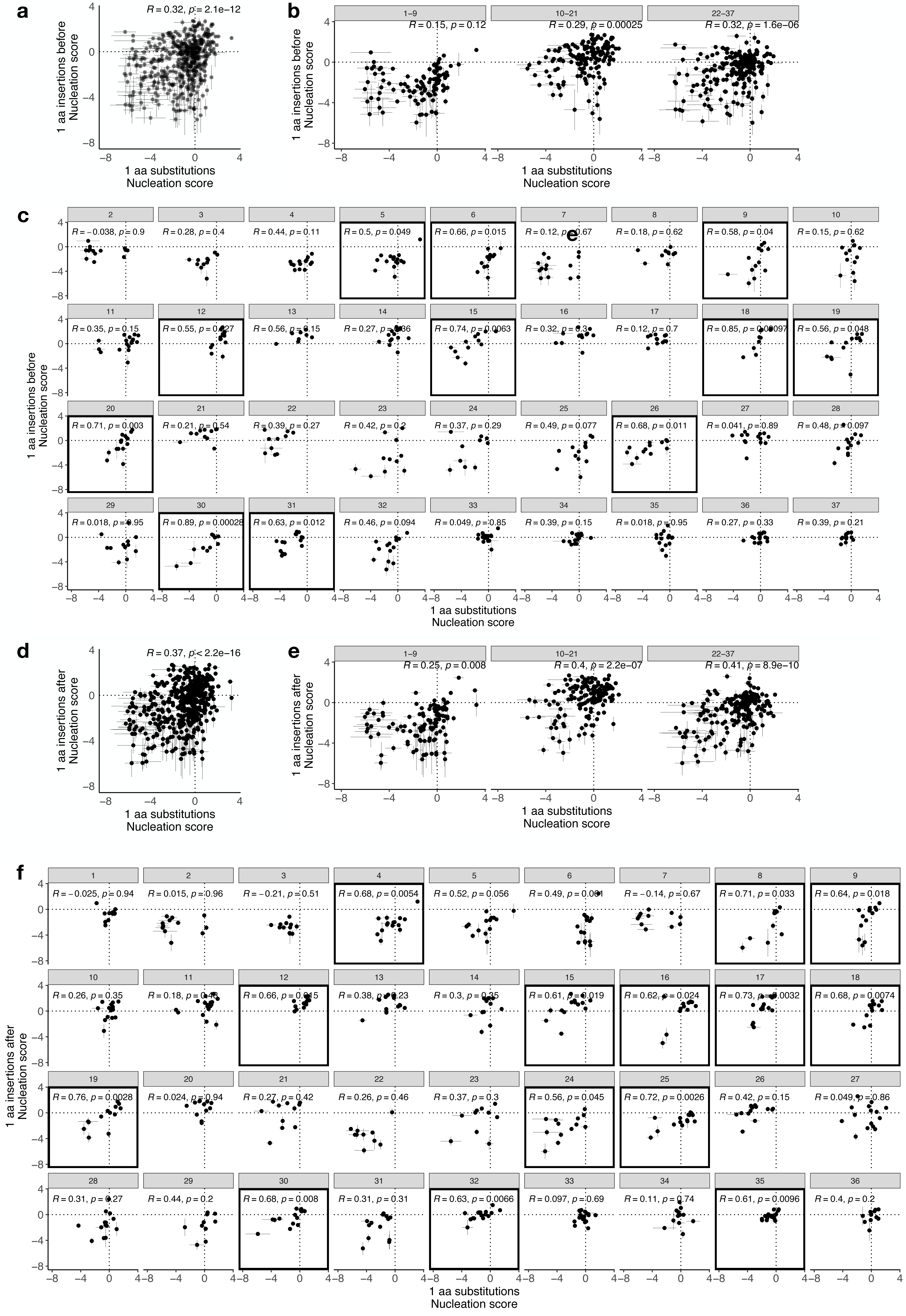
Comparison between the mutational effects of insertions and substitutions in IAPP. **a.** Correlation of nucleation scores resulting from single amino acid substitutions and insertions of the same amino acid before the substituted position. **b.** Same as a, but values are grouped by regions defined by hierarchical clustering and **c.** grouped by position. **d.** Correlation of nucleation scores of single amino acid substitutions and single amino acid insertions of the same amino acid inserted after a certain position. **e.** Correlation of nucleation scores of single amino acid substitutions and single amino acid insertions of the same amino acid inserted before a certain position grouped by regions defined by hierarchical clustering and **f.** grouped by position. Pearson correlations are indicated inside each correlation plot. Black squares highlight positions where correlations are significant in **c.** and **f**.

**Supplementary Figure 10.**
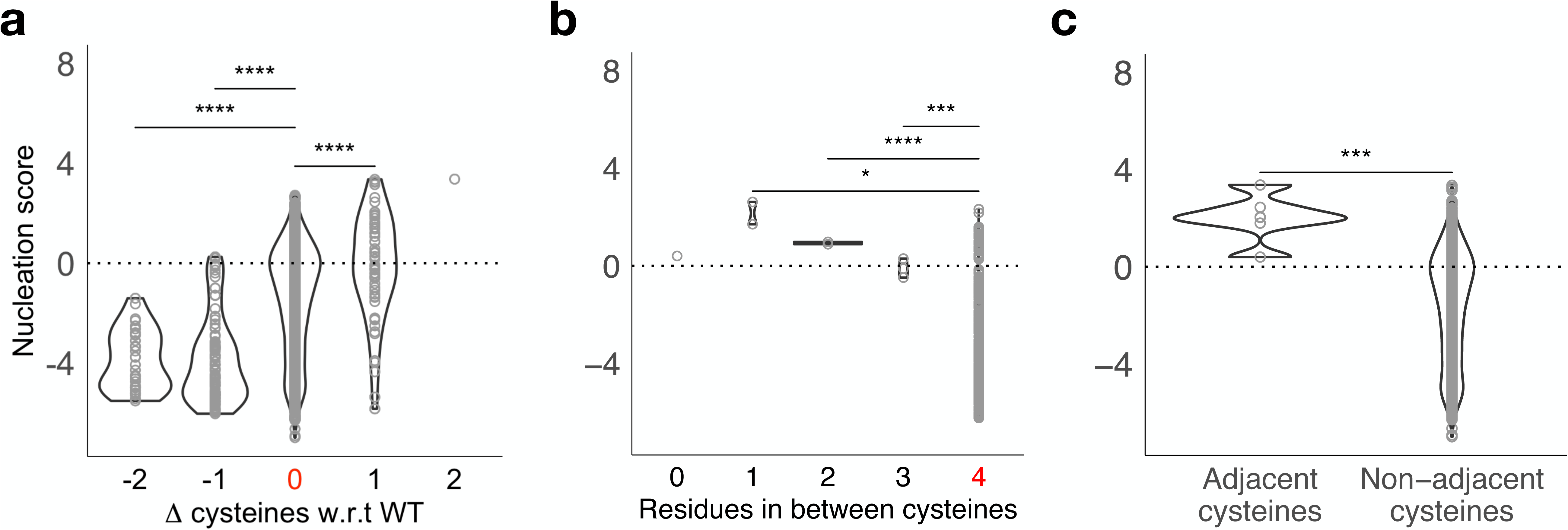
Mutational effect of cysteines in IAPP nucleation. **a.** Nucleation score distributions of IAPP variants (n = 1663) grouped by the total number of cysteines in their sequence. The x-axis indicates the difference in cysteine residues compared to the WT sequence (highlighted in red). **b.** Nucleation score distributions of IAPP deletions grouped by the distance between the 2 cysteine residues. Only deletions containing 2 cysteines in their sequence were considered for this plot. (n = 311). **c.** Nucleation scores of IAPP variants (n = 6) that have two adjacent cysteines in their sequence compared to the rest of IAPP variants.

**Supplementary Figure 11.**
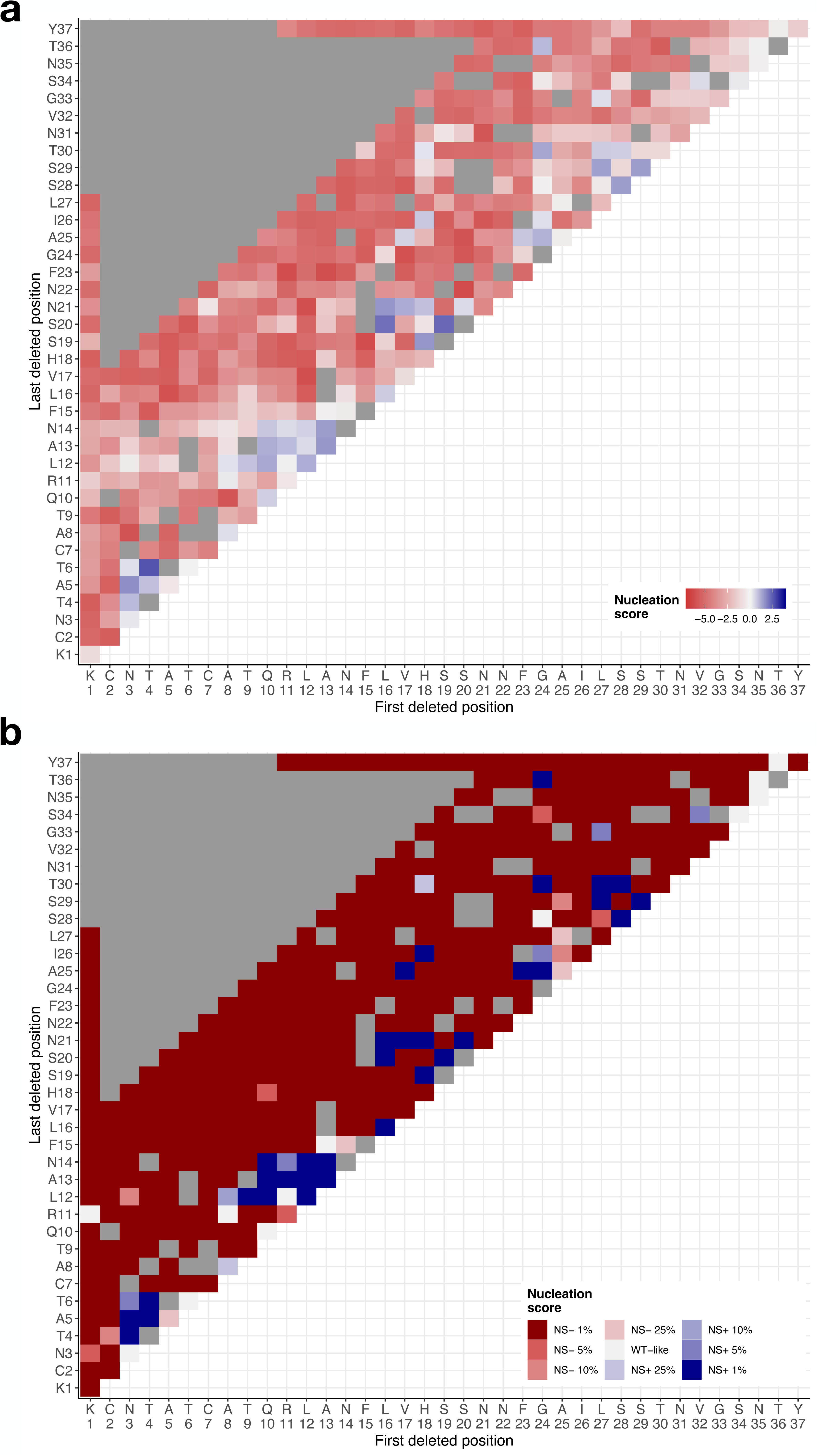
Multi amino acid deletions. Heatmap of **a.** nucleation scores and **b.** nucleation score FDR categories for IAPP single and multiple amino acid deletions. x-axis and y-axis indicate the first and the last residue deleted. Missing deletions are coloured in gray.

**Supplementary Figure 12.**
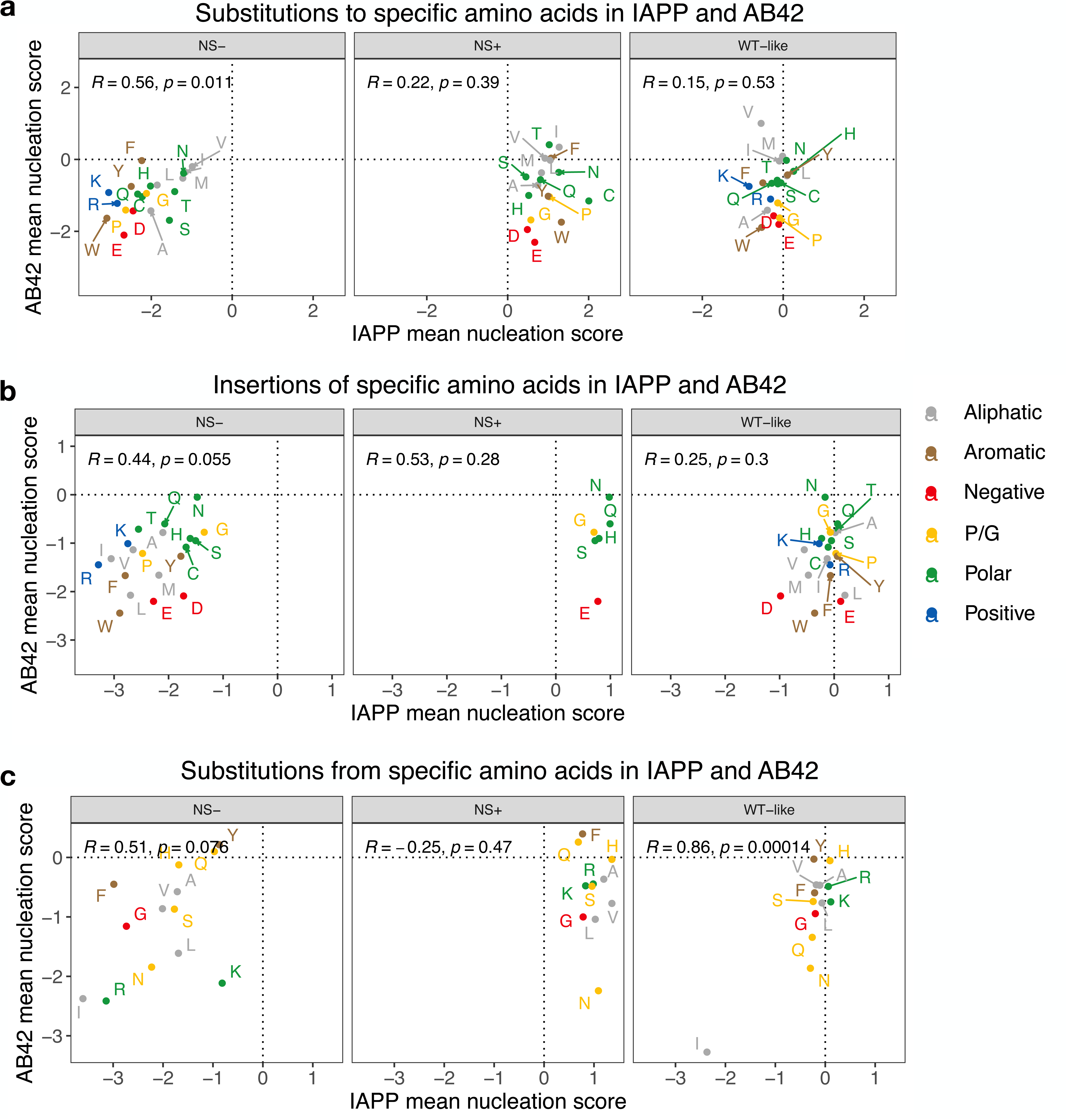
Correlation of single amino acid substitutions and insertions of IAPP and Aβ42 grouped by FDR categories. **a.** Correlation of the mean nucleation scores of substitutions to specific amino acids, **b.** insertions of to specific amino acids and **c. s**ubstitutions from specific amino acids in IAPP and Aβ42 grouped by their effect on nucleation (FDR = 0.1).

**Supplementary Figure 13.**
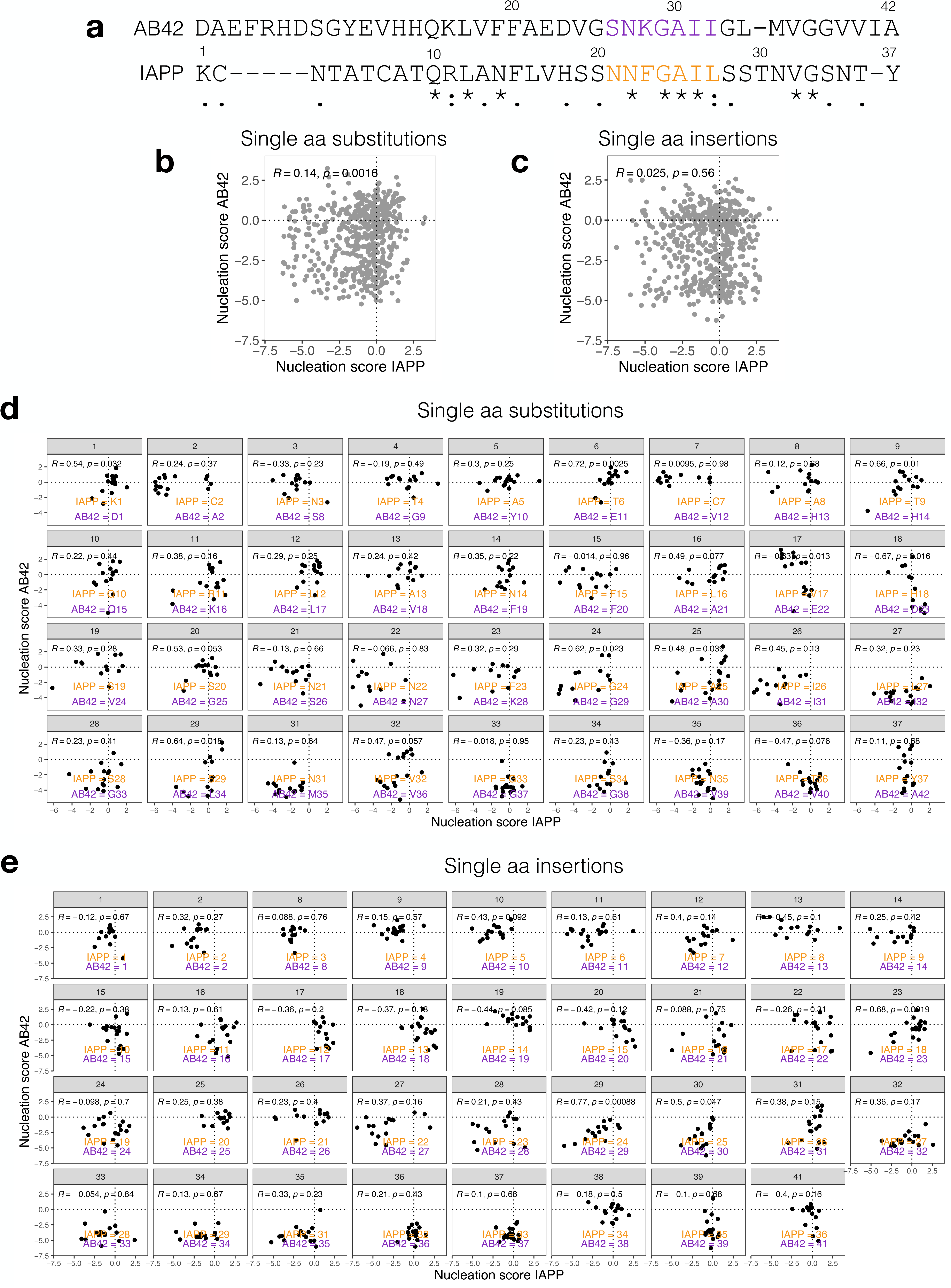
Correlation of single amino acid substitutions and insertions of IAPP and Aβ42 grouped by aligned position. **a.** Sequence alignment of IAPP and Aβ42 sequence by T-COFFEE^37^. WT positions of each protein are indicated above its sequence and gaps are indicated by “-”. Conservation scores are shown below the alignment: “*”, “:”, “.” indicate identical amino acids, conservative changes, and semi-conservative changes, respectively. **b.** Correlation of the nucleation scores of single amino acid substitutions and **c.** single amino acid insertions of IAPP and Aβ42 aligned by T-COFFEE. **d.** Correlation of the nucleation scores of single amino acid substitutions and **e.** single amino acid insertions of IAPP and Aβ42 grouped by T-COFFEE aligned position. WT amino acid and position are indicated in orange (IAPP) and purple (Aβ42).

**Supplementary Figure 14.**
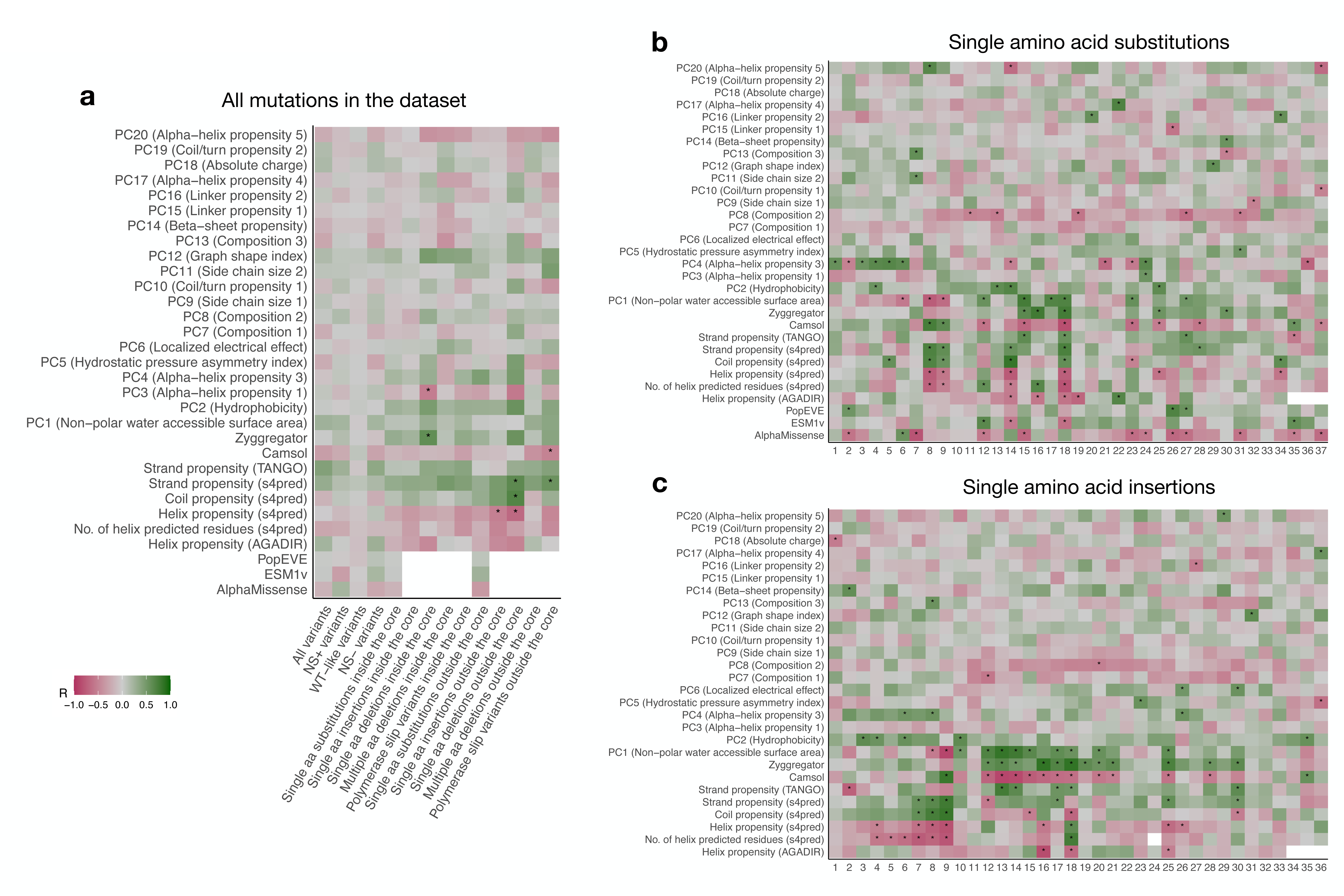
Comparison of nucleation scores and aggregation and variant effect predictors. **a.** Pearson correlation of nucleation scores with the predictions of secondary structure (Tango^47^, s4pred^48^), aggregation (Zyggregator^49^, Camsol^50^), amino acid physicochemical properties and variant effect predictors (PopEVE^51^, AlphaMissense^52^). IAPP variants are grouped by their nucleation effect or by the mutation and location type (inside core: residues 15-32), as indicated on the x-axis. Correlation coefficients are shown by colour, with “*” denoting correlations with p-values < 0.05 and |r| > 0.5. **b.** Correlation of nucleation scores and with predictor values, grouped per position in single amino acid substitutions. **c.** Correlation of nucleation scores and with predictor values, grouped per position in single amino acid insertions (after the position indicated in the x-axis).

**Supplementary Figure 15.**
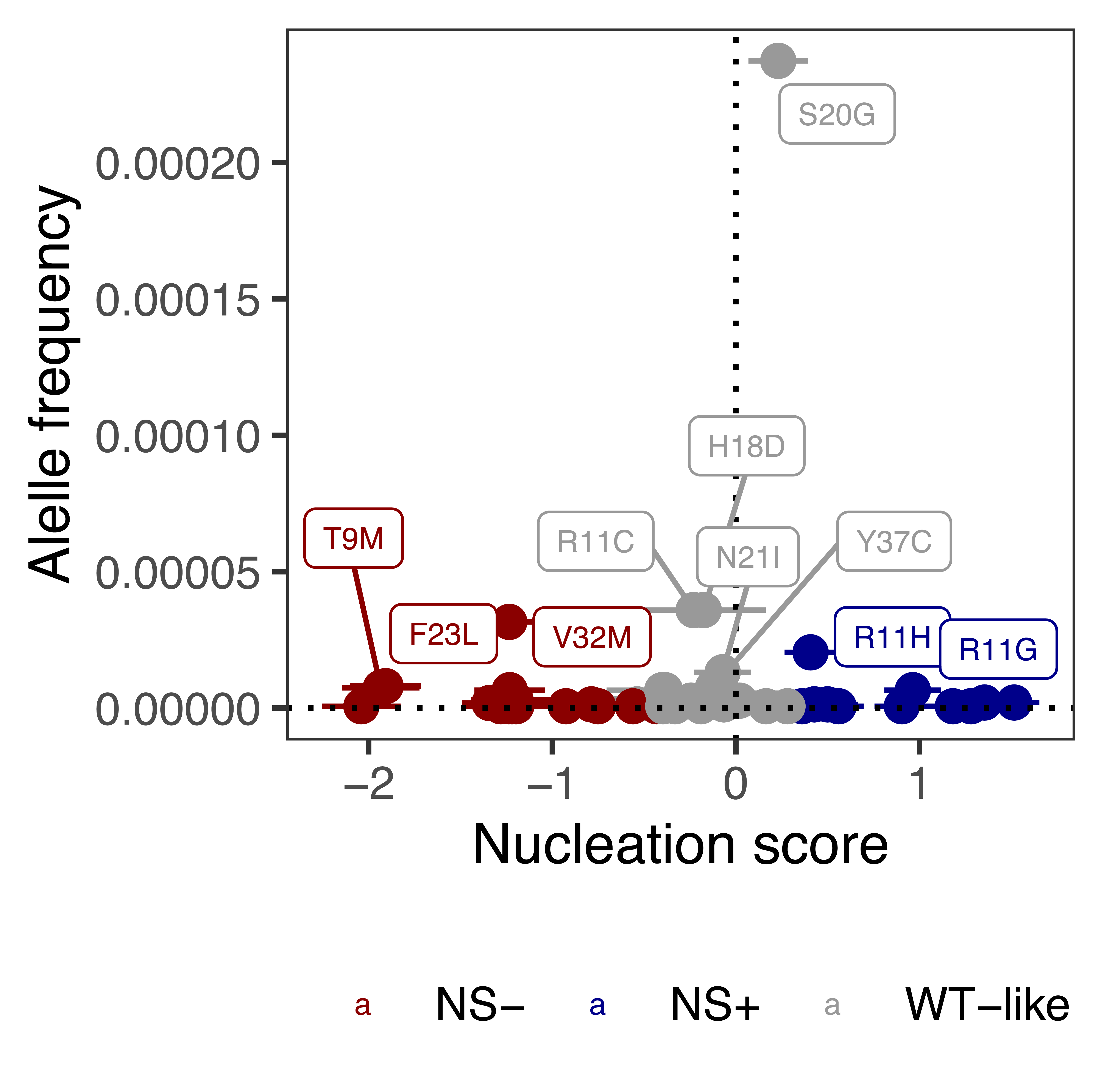
Allele frequency and nucleation scores. Comparison of gnomAD allele frequencies of IAPP variants and nucleation scores. Variants are colored by their FDR = 0.1 nucleation categories.

**Supplementary Table S1.** Impact on aggregation rates for IAPP variants for which these measurements could be retrieved from the literature.

**Supplementary Table S2.** IAPP nucleation scores obtained in this study, associated error estimates and their effect on nucleation (NS+, WT-like or NS-, FDR = 0.1).

**Supplementary Table S3.** List of oligonucleotides used in this study.

**Supplementary Table S4.** Burden, SKAT, Optimal Unified Tests (SKAT-O) and odds ratio for IAPP rare variant association analysis with diabetes and HbA1c levels. The different analyses were performed using either all SNPs found in the UK Biobank or by grouping them based on their effect on IAPP nucleation.

## Methods

### Library design

The designed library consists of 2000 unique IAPP variants and includes all possible single amino acid substitutions and stop codons at each position (n = 738), all single amino acid insertions at all positions (n = 685), all single amino acid deletions (n = 32), variants with bigger internal deletions with length ranging from 2 to 16 amino acids (n = 374), truncations of the N-terminal (n = 27) or the C-terminal (n = 27), 73 variants resulting from polymerase slippage, synonymous variants (n = 43) and the IAPP WT sequence.

### Plasmid library construction

The synthetic library was synthesized as an oligo pool by Twist Bioscience and with oligos consisting of IAPP variants region of 30 nt to 117 nt, flanked by 25 nt upstream and 21 nt downstream constant regions. 10 ng of the library were amplified by PCR (Q5 high-fidelity DNA polymerase, NEB) for 10 cycles with primers annealing to the constant regions (primers MB_01-02, Supplementary Table S3), following the manufacturer’s protocol. The product was treated with 2 ul of ExoSAP (Affymetrix) and then purified by column purification (MinElute PCR Purification Kit, Qiagen). The PCUP1-Sup35N plasmid^53^ was linearized by PCR (Q5 high-fidelity DNA polymerase, NEB) (primers MB_03-04, Supplementary Table S3). The product was treated with DpnI (ThermoFisher Scientific) and purified from a 1% agarose gel (QIAquick Gel Extraction Kit, Qiagen).

The library was ligated to 100 ng of the linear vector with a ratio of 5:1 (library:vector) by a Gibson approach with 3 h of incubation. The reaction product was dialyzed for 30 minutes on a membrane filter (MF-Millipore 0,025 um membrane, Merck) and transformed into 10-beta Electrocompetent *E.coli* (NEB), by electroporation with 2.0kV, 200 Ω, 25 uF (BioRad

GenePulser machine). Cells were recovered in SOC medium for 30 minutes and grown overnight in 50 ml of LB ampicillin medium. A number of cells were plated in LB ampicillin plates to calculate the transformation efficiency and confirm that each variant in the library would be represented at least 10 times. A total of 116.000 transformants were estimated. A total of 2 ml of overnight culture was harvested to obtain a miniprep of the IAPP library (NZYMiniprep kit, NZYTech).

### Yeast transformation

*Saccharomyces cerevisiae* [psi-][pin-] (MATa ade1-14 his3 leu2-3,112 lys2 trp1 ura3-52) strain was used in all experiments in this study. Yeast cells were transformed with the IAPP plasmid library in 3 independent biological replicates. An individual colony was grown overnight in 3 ml of YPDA medium at 30°C and 400 rcf. Cells were diluted to OD600 = 0.3 in 60 ml of YPDA medium and grown at 30°C and 400 rcf for 4h. Cells were harvested and split into 4 transformation tubes of 15 ml each. Each tube was treated as follows: cells were harvested at 400 rcf for 5 minutes and washed in 1 ml of YTB (100 mM LiOAc, 10 mM Tris pH 8, 1 mM EDTA). They were harvested again and resuspended in 72 ul YTB. 150 ng of the IAPP plasmid library, 8 μl of previously boiled ssDNA 10mg/uL (UltraPure, Thermo Scientific), 60 ul of DMSO, and 500 ul of YTB+PEG (100 mM LiOAc, 10 mM Tris pH 8, 1 mM EDTA pH 8, 40% PEG3350) were added to the cells. Heat shock was performed for 14 minutes at 42°C in a thermo block. Finally, cells were harvested and resuspended, and the 4 transformation tubes were pooled and added to a conical flask with 50 ml plasmid selection medium (-URA, 2% glucose). Cells were grown for 50 hr at 30°C and 400 rcf. Transformation efficiency was calculated for each biological replicate by plating an aliquot of cells in plasmid selection plates. A total of 345.000, 102.000 and 264.000 transformants were estimated for each biological replicate respectively, meaning that each variant in the library is represented at least 51 times. After 50h, the culture was diluted to OD600 = 0.05 in 50 ml plasmid selection medium and grown until OD = 0.8-1. Cells were harvested and stored at −80°C in 25% glycerol.

### Selection assays

*In vivo* selection assays were performed in 2 technical replicates for each biological replicate. For each technical replicate, cells were thawed from −80 °C in 50 ml plasmid selection medium at OD = 0.05 and grown until exponential for 15 h. At this stage, cells were harvested and resuspended at OD = 0.1 in 50 ml protein induction medium (-URA, 2% glucose, 100 µM Cu2SO4). After 24 h, input pellets were collected (10 ml per technical replicate) and stored at −20°C for later DNA extraction. 135 million cells/replicate were plated on -ADE -URA selection medium in 145cm2 plates (Nunc, Thermo Scientific). Plates were incubated at 30 °C for 6 days inside an incubator. Finally, colonies were scraped off the plates with PBS 1X and harvested by centrifugation to collect the output pellets, which were stored at −20 °C for later DNA extraction.

### DNA extraction and sequencing library preparation

One input and one output pellet for each technical and biological replicate (3 x 2 x 2 samples) were resuspended in 0.5 ml extraction buffer (2% Triton-X, 1% SDS, 100 mM NaCl, 10 mM Tris-HCl pH 8, 1 mM EDTA pH 8). They were then frozen for 10 minutes in an ethanol-dry ice bath and heated for 10 minutes at 62 °C. This cycle was repeated twice. 0.5 ml of phenol:chloroform:isoamyl (25:24:1 mixture, Thermo Scientific) was added together with glass beads (Sigma). Samples were vortexed for 10 min and centrifuged for 30 min at 20000 rcf. The aqueous phase was then transferred to a new tube, and mixed again with 0.5 ml of phenol:chloroform:isoamyl, vortexed, and centrifuged for 45 min at 20000 rcf. Next, the aqueous phase was transferred to another tube with 1:10 V 3 M NaOAc and 2.2 V cold ethanol 96% for DNA precipitation. After 30 min at −20 °C, samples were centrifuged and pellets were dried overnight. The following day, pellets were resuspended in 0.3 ml TE 1X buffer and treated with 10 µl RNAse A (Thermo Scientific) for 30 min at 37 °C. DNA was finally purified using 10 µl of silica beads (QIAEX II Gel Extraction Kit, Qiagen) and eluted in 30 µl elution buffer. Plasmid concentrations were measured by quantitative PCR with SYBR green (Merck) and primers annealing to the origin of the replication site of the PCUP1-Sup35N-IAPP plasmid at 58 °C for 40 cycles (primers MB_05-MB_06, Supplementary Table S3).

The library for high-throughput sequencing was prepared in a two-step PCR (Q5 high-fidelity DNA polymerase, NEB). In PCR1, 30 million molecules were amplified for 15 cycles with frame-shifted primers with homology to Illumina sequencing primers (primers MB_07-MB_20, Supplementary Table S3). The products were treated with ExoSAP (Affymetrix) and purified by column purification (MinElute PCR Purification Kit, Qiagen). They were then amplified for 10 cycles in PCR2 with Illumina-indexed primers (primers MB_21-MB_28, Supplementary Table S3). The six samples (3 inputs and 3 outputs) of each technical replicate were pooled together equimolarly and the final product was purified from a 2% agarose gel with 20 µl silica beads (QIAEX II Gel Extraction Kit, Qiagen).

The library was sent for 125 bp paired-end sequencing in an Illumina NextSeq500 sequencer at the CRG Genomics core facility. In total, >65 million paired-end reads were obtained, which correspond to 1.3 - 6.3 million per sample (i.e., input or output for a specific technical and biological replicate), representing >665x read coverage for each designed variant in the library.

### Individual variant testing

Selected IAPP single substitutions for individual testing were obtained by PCR linearization of the PCUP1-Sup35N-IAPP plasmid (Q5 high-fidelity DNA polymerase, NEB) with mutagenic primers (primers MB_28-MB_53, Supplementary Table S3). IAPP animal variants were obtained by ultramer amplification (primers MB_54-57, Supplementary Table S3) and Gibson assembly with the PCUP1-Sup35N-IAPP linearized plasmid (primers (primers MB_01-02, Supplementary Table S3). PCR products were treated with DpnI overnight and transformed into DH5 alpha-competent *E. coli.* Plasmids were purified by mini prep (NZYMiniprep kit, NZYTech) and transformed into yeast cells using one transformation tube of the transformation protocol described above. All constructions were verified by Sanger sequencing.

To determine the % growth in an adenine-lacking medium, yeast cells expressing individual variants were grown overnight in plasmid selection medium until exponential (-URA, 2% glucose). Protein expression then was induced by diluting the cells to OD 0.05 in protein induction medium (-URA, 2% glucose 100 µM Cu_2_SO_4_) and grown for 24 h. Cells were plated on -URA (control) and -ADE-URA (selection) plates in three independent replicates, and grown for 6 days at 30 °C. Growth in an adenine-lacking medium was calculated as the percentage of colonies in -ADE-URA relative to colonies in -URA.

### Data processing

FastQ files from paired-end sequencing of the IAPP library were processed using DiMSum^54^ (available at: https://github.com/lehner-lab/DiMSum), an R pipeline for analysing deep mutational scanning data. 5′ and 3′ constant regions were trimmed, allowing a maximum of 20% of mismatches relative to the reference sequence. Sequences with a Phred base quality score below 30 were discarded. Non-designed variants were also discarded for further analysis, as well as variants with fewer than 5 input reads in any of the replicates and variants resulting from one single nucleotide change with fewer than 70 input reads. Estimates from DiMSum were used to choose the filtering thresholds.

### Nucleation scores and error estimates

The DiMSum package (https://github.com/lehner-lab/DiMSum)^54^ was also used to calculate nucleation scores (NS) and their error estimates for each variant in each biological replicate as: Nucleation Score (NS) = ES_i_ - ES_WT_. Where, for a specific variant: ESi = log(F_i_ output) - log(F_i_ input) and for IAPP WT: ES_WT_ = log(F_WT_ output) - log(F_WT_ input). NSs for each variant were merged across biological replicates using error-weighted mean and centered to the mean of the NS of IAPP WT and synonymous variants. Sigma values were normalized to the interquartile range and variants with a normalized sigma value above a cut-off of 0.647 (¼ of the interquartile range) were excluded. Also, variants with sigmas higher than the threshold (due to their low output counts) that had high input counts (> 100 input counts in at least 1 replicate) were included in the dataset (n = 22). All NS and associated error estimates are available in Supplementary Table S2.

### Data analysis

#### Variants in the library

NS was obtained for 1663 unique IAPP variants, which were split into mutation classes: 599 single amino acid substitutions, 557 single amino acid insertions, 26 single amino acid deletions, 361 internal multi amino acid deletions, 59 variants resulting from polymerase slippage, 61 truncations (from either N-terminal or C-terminal), 35 synonymous variants and WT IAPP.

For single amino acid insertions, we assign the position of the inserted amino acid (e.g., an insertion between positions 1 and 2 is an insertion at position 2). Different mutations can result in the same coding sequence (e.g. Δ19-27, Δ20-28, and Δ21-29, which are all the same protein sequence: KCNTATCATQRLANFLVHSSTNVGSNTY). This is the case for single amino acid insertions, truncations and single and multi-amino acid deletions. They are only considered as one coding variant but considered multiple times for visualization or if the analysis is position-specific, in figures: Fig. 4 and Supplementary Figs. 6, 8, 9, 10 and 11).

We used Euclidean distance as the distance measurement to perform hierarchical clustering of the NS of single amino acid substitutions and insertions in a position-wise manner. Based on the clustering we defined 3 regions with different impact of the insertions: an N-terminal region (aa 1-9), a central region (aa 10-21) and a C-terminal region (aa 22-37).

#### Aggregation, amyloid nucleation and variant effect predictors

For the aggregation, solubility and secondary structure predictors^47,49,50^ (Tango, Zyggregator, Camsol and s4pred^48^), individual residue-level scores were summed to obtain a score for each IAPP variant and normalised by the sequence length. Alpha- and beta-propensity from Tango were used for correlation. For the “No. of helix predicted residues” value, residues predicted as helical in s4pred were counted for each IAPP variant and normalised by the sequence length.

For PopEVE^51^, we downloaded the pre-computed popEVE scores for IAPP from the PopEVE server and used the “pop.adjusted_EVE” and the “ESM1v” values for correlation. For AlphaMissense^52^, the pre-computed pathogenicity scores were obtained from the AlphaMissense server.

Principal components from a previously published PCA^55^ that reduced redundancy of amino acid physicochemical properties were also used for correlation.

#### Correlations of relative accessible surface area and nucleation scores

Relative accessible surface area (ASA) of each residue in IAPP WT PDB structures (PDB IDs: 7M61, 7M62, 7M64, 7M65, 6Y1A, 6ZRF, 8R4I and 6VW2) were obtained using DSSP package from BioPython^56^ and correlated to the mean NS of each IAPP position. Residues are classified as buried if their relative ASA is < 0.25^57^.

#### Comparison of gnomAD and UK BioBank allele frequencies and nucleation scores

We extracted allele frequencies of IAPP variants (ENST00000240652.8, NM_000415.3) in the human population from gnomAD v4.1.0 ^38^.

Self-reported diabetes status, HbA1c levels, age, sex and body mass index were extracted for 501,252 UK Biobank participants, including 192 who carried a non-synonymous IAPP variant. Participants from the UK Biobank were classified into three different groups based on their HbA1c levels, according to the WHO guidelines for diabetes diagnosis: (i) no-risk: participants with HbA1c levels lower than 42 mmol/mol (n = 427269), (ii) pre-diabetes: participants with HbA1c levels between 42 and 48 mmol/mol (n = 21440) and (iii) diabetes: participants with HbA1c levels greater than 48 mmol/mol (n = 17448). No-risk and pre-diabetes participants were considered as controls and diabetes participants were considered as cases. Participants with HbA1c levels outside the analytical range of the instrument (15 - 184 mmol/mol) were excluded from the analysis. In parallel, participants were categorised on the basis of their response to the question: “Has a doctor ever told you that you have diabetes?” during their first visit as follows: (a) diabetes: participants that answered “YES” (n = 26386) and (b) no diabetes: participants that answered “NO” (n = 473128). To assess the association of IAPP variants and diabetes diagnosis, participants in (a) and (b) were considered cases and controls, respectively. Participants who didn’t select either options were excluded from the analysis. Allele frequencies for each group were retrieved from the UK Biobank for comparison to nucleation scores.

Association between IAPP variants and diabetes diagnosis or HbA1c levels were performed using the SKAT R package^58^. Age, sex and body mass index were used as covariates. Different SKAT analyses were performed using either all IAPP variants present in the UK Biobank or grouping IAPP variants depending on their effect in nucleation (NS+, WT-like and NS-). Burden, SKAT and SKAT-O tests p-values can be found in Supplementary Table S4. Odds ratios were calculated using a multivariable logistic regression model using Firth’s penalized likelihood method adjusting for age, sex and body mass index. IAPP variants were concatenated depending on their effect in nucleation (NS+, WT-like and NS-).

### Statistics and reproducibility

Based on the transformation efficiency, each variant in the designed libraries (n = 2000) is expected to be represented at least 10x at each step in the selection experiments and library preparation. In terms of sequencing, reads that did not pass the QC filters using the DiMSum package were excluded (https://github.com/lehner-lab/DiMSum). The experiments were not randomized. The investigators were not blinded to allocation during experiments and outcome assessment.

## Data availability

Raw sequencing data and the processed data table are deposited in NCBI’s Gene Expression Omnibus (GEO) as GSE281555. The processed data are provided in the Supplementary Table 2. The coordinates for the PDB structures used in the study were obtained with accession: 7M61, 7M62, 7M64, 7M65, 6Y1A, 6ZRF, 8R4I, 6VW2, 8AWT, 8AZ0, 8AZ1, 8AZ2, 8AZ3, 8AZ4, 8AZ5, 8AZ6, 8AZ7, 6ZRR, 6ZRQ, 7Q4M. Source data are provided with this paper.

## Acknowledgments

M.B. was supported by the fellowship “Ayudas para contratos predoctorales para la formación de doctores 2019” (PRE2019-088300) from the Spanish Ministry of Science, Innovation and Universities. Work in the lab of B.B. is supported by the la Caixa Research Foundation project ‘DeepAmyloids’ (LCF/PR/HR21/52410004), by the Spanish Ministry of Science, Innovation and Universities (PID2021-127761OB-I00 and RYC2020-028861-I, funded by MCIN/AEI/10.13039/501100011033, “ERDF A way of making Europe” and “ESF Investing in your future”). and by the European Union (ERC Consolidator, Glam-MAP, 101125484). Views and opinions expressed are however those of the author(s) only and do not necessarily reflect those of the European Union or the European Research Council. Neither the European Union nor the granting authority can be held responsible for them. IBEC is a member of the CERCA Program/Generalitat de Catalunya. This work uses data provided by patients and collected by the NHS as part of their care and support. This research has been conducted using the UK Biobank under the Application Number 441451 and in accordance with the UK Biobank Ethics and Governance Framework. UK Biobank data are publicly available by request from https://www.ukbiobank.ac.uk. We thank the Chernoff lab for providing strains and plasmids and the CRG Genomics core technology for sequencing. We thank Prof. John Perry, Dr. Yajie Zhao, Prof. Ben Lehner, Carla Folgado and the B.B. lab members for discussing our data.

## Notes

https://www.ncbi.nlm.nih.gov/geo/

### Competing Interest Statement

The authors have declared no competing interest.

